# Global kinetic model of lipid-induced *α*-synuclein aggregation and its inhibition by small molecules

**DOI:** 10.1101/2024.11.07.622437

**Authors:** Alisdair Stevenson, Roxine Staats, Alexander J. Dear, David Voderholzer, Georg Meisl, Raphael Guido, Céline Galvagnion, Alexander K. Buell, Tuomas P.J. Knowles, Michele Vendruscolo, Thomas C.T. Michaels

## Abstract

The aggregation of *α*-synuclein into amyloid fibrils is a hallmark of Parkinson’s disease. This process has been shown to directly involve interactions between proteins and lipid surfaces when the latter are present. Despite this importance, the molecular mechanisms of lipid-induced amyloid aggregation have remained largely elusive. Here, we present a global kinetic model to describe lipid-induced amyloid aggregation of *α*-synuclein. Using this framework we find that *α*-synuclein fibrils form via a two-step primary nucleation mechanism and that lipid molecules are directly involved in both the nucleation and fibril elongation steps, giving rise to lipid-protein coaggregates. To illustrate the applicability of this kinetic approach to drug discovery, we identify the mechanism of action of squalamine, a known inhibitor of α-synuclein aggregation, finding that this small molecule reduces the rate of lipid-dependent primary nucleation. Our work will likely guide the rational design of *α*-synuclein aggregation inhibitors.

**Significance Statement:** Amyloid aggregation is a hallmark of a diverse range of diseases including Parkinson’s Disease, where the protein *α*-synuclein is a major con-stituent of proteinaceous deposits found in patients. It is well established that interactions between *α*-synuclein and lipids modulate aggregation. However, the molecular mechanisms driving this lipid-induced aggregation have remained largely elusive, which has frustrated so far the discovery of drugs that prevent lipid-induced aggregation. In this work, we present the first global kinetic model describing lipid-induced aggregation and by integrating this theoretical framework with *in vitro* experimental data of lipid-induced *α*-synuclein aggregation, we reveal the role of lipid membranes in the aggregation process and uncover the mechanism by which small molecule inhibitors interfere with this process.

---

Parkinson’s disease (PD) is the second most common neurological ageing-related pathology. It is estimated that over 10 million people currently have PD worldwide and, owing to the ageing population, this number is expected to increase to over 17 million by 2040 (1). Therefore, there is a growing urgency to understand the mechanisms of PD pathogenesis and to develop effective therapeutic treatments. *α*-synuclein, a small (14kDa) intrinsically disorded protein, is implicated in both familial and sporadic PD as a hallmark of these pathologies is abnormal proteinaceous deposits, known as Lewy bodies, which consist of amyloid fibrils of *α*-synuclein and other components, including lipids (2–6). Identifying potential therapeutic compounds that inhibit the aggregation is a significant challenge given the complexity of the overall aggregation reaction with multiple microscopic steps, that are all potential targets (7, 8). Therefore, without a detailed understanding of the mechanistic pathway of aggregation and inhibition, it is challenging to accurately interpret and utilize *in vitro* data to aid therapeutic development.

At neutral pH, *α*-synuclein is relatively resistant to aggregation, likely due to slow fibril nucleation (9, 10). To overcome this slow rate, aggregation typically requires stirring (11), addition of pre-formed fibril seeds (9, 10), lowering pH (9, 12) or increasing salt concentrations (13), where changes to the pH and salt concentrations likely induce aggregation via minimizing the electrostatic repulsion between aggregating proteins (14). However, the N-terminal region and part of the non-amyloid core of *α*-synuclein can bind to lipid surfaces and these interactions may form part of the *in vivo* function and pathology of *α*-synuclein (3, 4, 6, 15–17). Notably, the interactions between *α*-synuclein and lipids have been shown to modulate aggregation (16–24).

From a structural perspective, *α*-synuclein fibrils formed in the presence of lipids are distinct from those formed without lipids (9, 21, 23). Pure *α*-synuclein fibrils are long, relatively straight structures (Fig. 1a). Fibrils formed in the presence of lipids appear more flexible in microscopy images (Fig. 1b) and are coaggregates of lipids and proteins (21, 23–28). At low lipid-to-protein ratios, lipids are closely associated with the fibril structure, while at high lipid-to-protein ratio values, excess lipids that are not incorporated into lipid-protein coaggregates can form vesicles that adsorb along the length of fibrils in some cases (25). In the structural conversion of soluble *α*-synuclein monomers to fibrils, the initial formation of intermediate oligomer species is of particular interest in PD pathogenesis as they interact extensively with lipid membranes and have been identified as a key cytotoxic species in PD (29). The toxicity of oligomers is thought to arise from the disruption they cause to functional lipid membranes that have been observed in synthetic systems (20, 29, 30), as well as for physiological lipid membranes derived from synaptic vesicles (28), and these events precede cellular stress and death *in vitro* (31). Finally, *α*-synuclein has specific interactions with the mitochondrial lipid, cardiolipin (32), and coaggregates of *α*-synuclein and cardiolipin have been observed in neuronal cells *in vitro* (26). The mitochondrial membrane disruption caused by the incorporation of functional lipids into coaggregates has been proposed as a mechanism of neuronal toxicity (26, 32), where mitochondrial damage precedes cell death, with oligomers specifically identified to have mitotoxic effects (33).

**Fig. 1.**
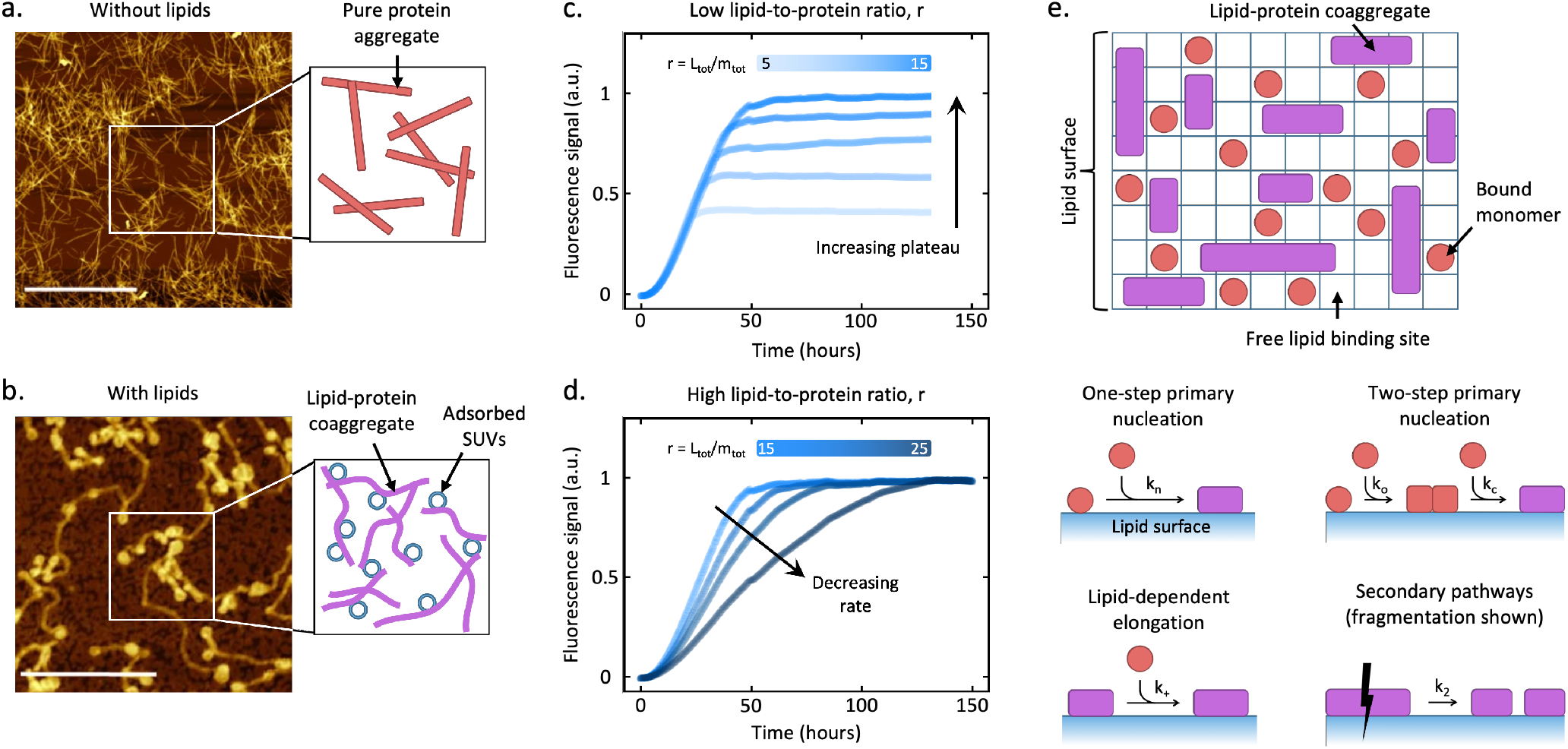
Interaction of *α*-synuclein with lipid surfaces affects *in vitro* aggregation kinetics and the kinetic model describing lipid-induced amyloid aggregation developed in this work. **(a)** Atomic force microscopy (AFM) image of *α*-synuclein fibrils formed in the absence of lipids, adapted from Galvagnion et al. (21), with a schematic representation of the qualitative features of the system. **(b)** AFM image of *α*-synuclein fibrils formed in the presence of lipids, adapted from Galvagnion et al. (21), with a schematic representation of the qualitative features of the system. **(c)** Experimental measurements of the *in vitro* kinetics of lipid-induced *α*-synuclein aggregation under varying initial lipid concentrations (constant *m*_tot_ = 20 *μ*M and *L*_tot_ = 100, 150, 200, 250, 300 *μM*). **(d)** Experimental measurements of the kinetics of lipid-induced *α*-synuclein aggregation under varying initial lipid concentrations (constant *m*_tot_ = 20 *μ*M and *L*_tot_ = 350, 400, 500 *μM*). **(e)** Top panel: Schematic representation of monomer and fibril surface coverage of a vesicle. Bottom panel: Elementary reaction mechanisms of lipid-induced aggregation considered in this study.

From an aggregation kinetics perspective, the role of the interaction between *α*-synuclein and lipids in amyloid aggregation has been investigated extensively *in vitro* (21–23). Vesicles of the phospholipid DMPS (1,2-dimyristoyl-sn-glycero-3-phospho-L-serine) have been shown to modulate *α*-synuclein aggregation either by accelerating or inhibiting aggregation depending on the lipid-to-protein ratio, highlighting the complex effects of the lipid-protein interactions (21). Similar effects have been observed with other synthetic lipid systems such as DLPS (1,2-dilauroyl-sn-glycero-3-phospho-L-serine) (34, 35), as well as for physiological lipids from synaptic vesicles isolated from rodent brains (28). At low lipid-to-protein ratios, the presence of lipids accelerates *α*-synuclein fibril formation by approximately 2-3 orders of magnitude compared to the absence of lipids and the final yield of aggregates increases with lipid concentration (Fig. 1c) (21). Increasing the lipid-to-protein ratio above a critical value, however, decelerates the aggregation process while not affecting the final yield of aggregates (Fig. 1d) (21). Chemical kinetics has proven to be a powerful mechanistic tool for analysing kinetic traces of protein aggregation, transforming our understanding of the molecular mechanisms relevant to filamentous protein aggregation for systems ranging from Amyloid-*β* to prions (7, 8, 36–43), as well as to *α*-synuclein under different experimental conditions and without lipids (9, 12). These chemical kinetics models of aggregation, however, cannot capture the unique features of lipid-induced aggregation since they model aggregates consisting only of protein, rather than lipid-protein coaggregates, and therefore predict, for example, that all protein monomers are converted into aggregates at the end of the reaction (7, 8, 36–43). Theoretical studies of lipid-induced *α*-synuclein aggregation so far have focused on describing the early-time aggregation kinetics (21), the regime of low lipid-to-protein ratios (44), or when lipid surfaces sequester protein monomers but are otherwise not involved in the aggregation reaction (45). Therefore, we currently lack a globally applicable chemical kinetics framework capable of quantitatively describing lipid-induced amyloid aggregation.

To overcome this challenge, we present a generalizable theoretical model of lipid-induced amyloid aggregation kinetics spanning different lipid-to-protein ratio regimes and derive analytical self-consistent integrated rate laws that can be used to achieve accurate global fits of experimental data and quantify the associated rate parameters for the formation and propagation of fibrils. We apply this framework to *in vitro α*-synuclein aggregation in the presence of DMPS to reveal the mechanistic pathway for this lipid-induced aggregation process and quantify the relevant rate parameters. The mechanistic insight into the lipid-induced aggregation of *α*-synuclein from our model enabled the informed analysis of the effect of a known inhibitor, squalamine, on the aggregation kinetics to elucidate the mechanism of inhibition and quantify the reduction of relevant rate parameters.

## Results

### Kinetic modelling of lipid-induced protein aggregation

The general approach to mathematically modelling protein aggregation kinetics in the absence of lipids uses differential equations to describe the time evolution of aggregate concentrations in terms of the different microscopic aggregation events (7, 36, 37, 40–43). We adapt this formalism to describe protein aggregation on lipid surfaces (see Fig. 1e for schematic representations). By analogy to surface reaction kinetics (see Supplementary Material Sec. S3 for an introduction) and since aggregation involves interactions with a lipid surface, we consider the surface coverage of protein monomers and aggregates as key quantities in our model instead of absolute concentrations. The surface coverage of protein monomers is defined as *θ*_*m*_ = *m*_b_*β/L*_tot_ (46, 47), where *m*_b_ is the concentration of protein bound to the lipid surface, *L*_tot_ is the total concentration of lipids (surface binding sites) and *β* = 28.2 *±* 0.8, measured by Galvagnion et al., is the average number of lipids that each protein monomer binds to when vesicles are fully covered by protein (21). For the fibrils, we distinguish between their number concentration *P* (moles of fibrils per unit volume) and mass concentration *M* (moles of aggregated protein monomers per unit volume) and introduce the respective surface coverage terms as *θ*_*P*_ = *P/L*_tot_ and *θ*_*M*_ = *Mα/L*_tot_, where *α* is the stoichiometry of lipid to protein in fibrils. We then derived systems of differential equations for the time evolution of the surface coverage of monomeric and fibrillar proteins in terms of different microscopic aggregation events (see Methods section for explicit models). Using self-consistent methods (40), we derived analytical solutions for the resulting kinetics which we used to fit experimental measurements of *α*-synuclein aggregation and test different mechanistic scenarios.

### Lipid-induced aggregation exhibits biphasic thermodynamic behavior with lipid-to-protein ratio

Using our kinetic model, we first sought to understand how aggregation is affected by the presence of a lipid surface when varying the lipid-to-protein ratio, *r* = *L*_tot_*/m*_tot_. The predicted kinetic traces display a biphasic thermodynamic behavior, where the reaction is either lipid- (Fig. 2a left panel) or protein monomer-limited (Fig. 2a right panel). We distinguish the situations when *r* is varied by (i) varying *L*_tot_ while keeping *m*_tot_ constant or (ii) varying *m*_tot_ while keeping *L*_tot_ constant. In the first case, at low *r* values the total aggregate concentration yield is limited by lipid concentration and a significant fraction of the total protein concentration remains unreacted in monomer form when the aggregation reaction stops (Fig. 2b). Increasing *r* causes the plateau concentration to increase as *M*(∞) = *L*_tot_*/α* and the fraction of unreacted protein monomers at the end of the reaction is reduced. This behavior is in distinct contrast to aggregation kinetics without lipids, where the total protein mass will usually be converted nearly completely into fibril form, i.e. *M*(∞) = *m*_tot_, and there is no shift of plateau, assuming the rate of protein monomer dissociation from fibril ends is negligible. If we increase *r* beyond a critical value, *r*^∗^, the lipids are now available in excess relative to the protein concentration. In this case, the plateau of fibril mass concentration corresponds to the full conversion of protein from monomeric to fibril form, *M*(∞) = *m*_tot_ (assuming fibril dissociation is negligible), and no longer depends on *L*_tot_. Interestingly, increasing *r* in the protein-limited regime causes a reduction in the overall rate of aggregation. In our model, this behavior emerges because the key quantity that determines the rate of lipid-induced fibril proliferation is protein monomer surface coverage rather than concentration. As we increase *r*, we dilute the protein monomer surface coverage due to the higher number of available lipid binding sites. As a result, aggregation slows down. Although our model assumes non-cooperative binding of protein monomers to the lipid surface, the same deceleration of aggregation is observed with a model of cooperative binding (see Supplementary Material Sec. S5 for details) (17).

**Fig. 2.**
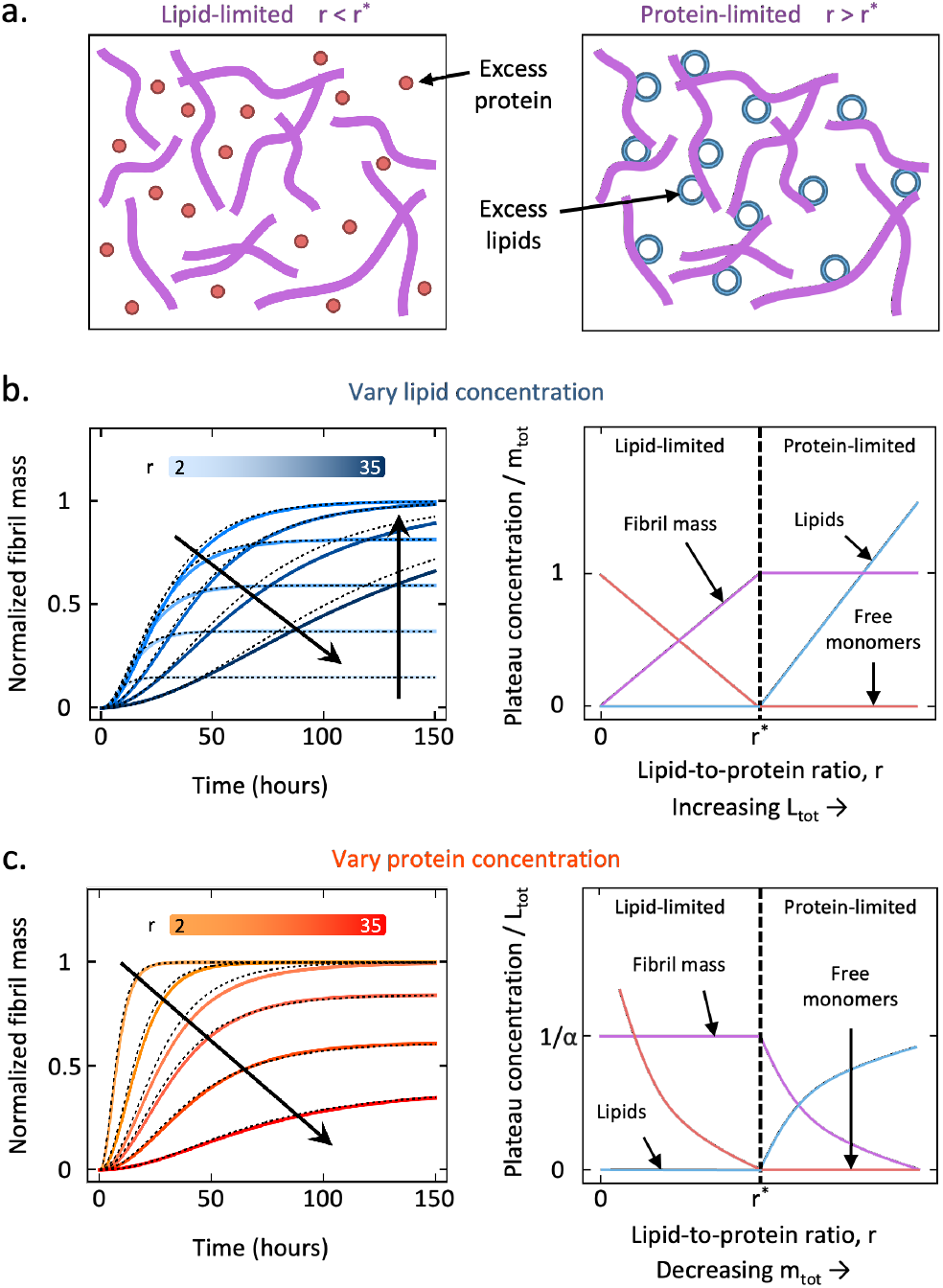
Predicted biphasic kinetics and steady-state behaviors from the kinetic model. **(a)** Left panel: Schematic representation of the system at steady-state when the lipid-to-protein ratio, *r* = *L*_tot_*/m*_tot_, is below the critical value *r*^∗^. In this case, fibril formation is limited by the lipid concentration and unreacted protein monomers remain in solution. Right panel: Schematic representation of the system at steady-state when *r > r*^∗^. In this case, aggregation is limited by the protein monomer concentration, resulting in all monomeric proteins converting to fibril form and an excess of lipids remaining. **(b)** Left panel: Dynamics of lipid-induced aggregation for increasing *r* values (increasing *L*_tot_). The solid lines indicate the numerical solution to the differential equations describing the one-step primary nucleation model and the dashed lines are the self-consistent analytical solution to this model. Right panel: The effect of increasing the lipid-to-protein ratio, *r*, by increasing *L*_tot_ on the free protein monomer, aggregate mass and lipid plateau concentrations. **(c)** Left panel: Dynamics of lipid-induced aggregation for increasing *r* values (decreasing *m*_tot_) where the solid lines are predictions from the differential equations and the dashed lines are from the self-consistent solution. Right panel: The effect of increasing the lipid-to-protein ratio, *r*, by decreasing *m*_tot_ on the free protein monomer, aggregate mass and lipid plateau concentrations.

A similar biphasic thermodynamic behavior is observed when varying *m*_tot_ while keeping *L*_tot_ constant (Fig. 2c). At low *r* values, the yield of the reaction is lipid-limited and as the lipid concentration is constant, the fibril plateau concentration does not change with *r*. As we increase *r* beyond *r*^∗^, the yield of the reaction is now protein-limited and all protein monomers will be converted into fibrils, which causes the plateau and rate of aggregation to decrease with increasing *r*. Our model thus captures the biphasic thermodynamic behavior experimentally observed from *in vitro* lipid-induced *α*-synuclein aggregation (Fig. 1c and d) and rationalize why the presence of lipids can either enhance or inhibit amyloid formation depending on the value of *r* (18, 19, 21, 30).

### Analysis of experimental data for lipid-induced α-synuclein aggregation

We next sought to use our model to interpret *in vitro* measurements of *α*-synuclein aggregation in the presence of small unilamellar vesicles (SUVs) of DMPS, which are significantly smaller than the length of the fibrils produced by the end of the experiment. Lipid-induced *α*-synuclein aggregation was measured by ThT fluorescence where either the initial lipid concentration was varied, with protein concentration constant (Fig. 3a) or the initial protein concentration was varied, with lipid concentration constant (Fig. 3b). Using our model we simulated different mechanistic scenarios to assess the level of quantitative agreement between the experimental data and theory using a global fitting procedure. In the simplest scenario, fibrils are formed via one-step primary nucleation followed by elongation with both steps involving protein and lipid molecules (see Eq. 2). Nucleation can involve variable combinations of proteins and lipids depending on the relevant reaction orders. Figs. 3a and b show that the best fit of this model to the data is not able to capture all features of the data, particularly when protein concentration was varied. To identify the missing mechanism necessary for the model, we considered a two-step primary nucleation process where, generally, nucleation proceeds via intermediate oligomers that need to convert into elongation-competent mature fibrils (see Eq. Eq. (6)) as was previously suggested for lipid-induced *α*-synuclein aggregation (21). Two-step primary nucleation occurs in many amyloid forming systems (48–50), and in nucleation processes more generally (43, 51–53). This model shows significantly improved agreement with the data (Figs. 3c, d) with a summary of the fitting parameters shown in Supplementary Material Table S5. The necessity to include a two-step nucleation mechanism to describe our data is most clearly evidenced when analyzing the early-time behavior of the aggregate mass concentration. In particular, mathematical analysis of the theoretical model with one-step nucleation predicts the scaling of the aggregate mass with time to be of quadratic form *t*^2^ at the early stages of the aggregation reaction. With the introduction of a two-step nucleation mechanism, the early-time behavior is described by a higher power, *M* ∼*t*^*n*^ (2 ≤*n <* 3) (21), which agrees with the experimental data (Supplementary Material Fig. S1). One key fitting parameter in our model is the stoichiometry of lipid to protein in fibrils, *α*, which has previously been found to exhibit a dependence on lipid concentration, with the value for the most thermodynamically stable state being around *α* = 15, but reducing down to as low as *α* = 10 when the aggregation reaction is lipid-limited (44). From our global fitting analysis, we estimate the average number of lipids incorporated per protein monomer into fibrils to be approximately *α* = 12.6 *±* 1.9 across the experiments analyzed here (see Table S7 for a summary). Therefore, fewer lipids will be able to bind to proteins in fibril form compared to the monomeric state (*β* = 28.2 *±* 0.8). We also considered the possibility of secondary mechanisms, including fragmentation, being present in the aggregation pathway with the best fit of the kinetic model to the data shown in Supplementary Material Fig. S3. While this model can describe the experimental data, it requires significantly more fitting parameters when compared to the two-step primary nucleation model.

**Fig. 3.**
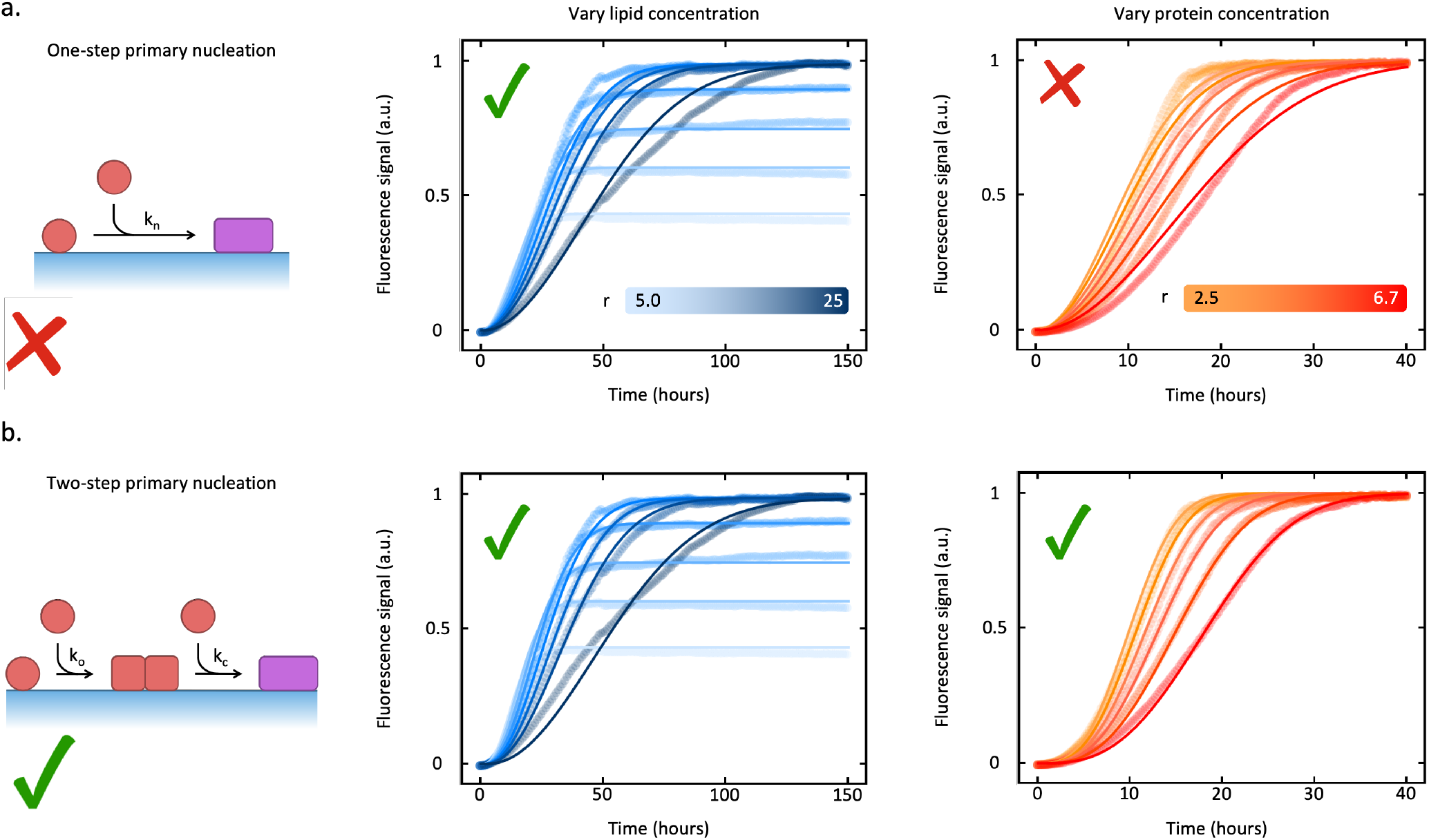
Kinetic analysis of *in vitro α*-synuclein aggregation in the presence of lipid vesicles reveals the mechanistic pathway. Experimental measurements of the time evolution of *in vitro* lipid-induced *α*-synuclein aggregation, under varying initial lipid (middle panels) and *α*-synuclein monomer (right panels) concentrations, with best fits to the self-consistent analytical solutions describing the chemical kinetics model with single-step primary nucleation model **(a)**, or with two-step primary nucleation model **(b)** using Eq. Eq. (9). The middle panels show aggregation curves monitoring the kinetics of fibril mass formation using ThT-fluorescence with constant *m*_tot_ = 20 *μ*M and varying *L*_tot_ = 100, 150, 200, 250, 300, 350, 400, 500 *μM*. Fit parameters summarized in Supplementary Material Table S3. The right panels show aggregation curves with constant *L*_tot_ = 100 *μ*M and varying *m*_tot_ = 15, 20, 25, 30, 35, 40 *μM*. Fit parameters summarized in Supplementary Material Table S5. Reaction orders were taken from the values determined from *t*_1*/*2_ scaling analysis (Fig. 4).

### Half-time scaling reveals reaction orders

To gain further mechanistic insights into the two-step nucleation process and to determine the reaction orders involved, we have employed a methodology based on the scaling of the reaction half-time, *t*_1*/*2_. The half-time is the time required for the fibril mass concentration to achieve half its maximum value and is a crucial metric in the study of amyloid aggregation (7, 40–43). Prior investigations into aggregation in the absence of lipids revealed that *t*_1*/*2_ follows a scaling relation 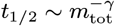 (7, 40–43) with monomer concentration *m*_tot_. The scaling exponent *γ* depends on the reaction orders of the prevailing nucleation process and has thus proven useful in deducing the dominant nucleation mechanisms during fibril formation in systems like amyloid-*β* (7, 43).

In the context of lipid-induced aggregation, we can vary the initial concentrations of both the protein, *m*_tot_, and lipids, *L*_tot_, giving rise to richer scaling behavior of *t*_1*/*2_, which we capture by means of two scaling exponents *γ*_*p*_ and *γ*_*L*_ describing the dependence of *t*_1*/*2_ on *m*_tot_ and *L*_tot_ respectively. We identify four distinct regimes of *t*_1*/*2_ scaling, depending on whether *r < r*^∗^ or *r > r*^∗^ and if the timescale of the conversion of oligomers to fibrils is slow or fast compared to the characteristic timescale of aggregation, *t*_1*/*2_ (Fig. 4). At lower lipid-to-protein ratios than considered in this work, the incorporation of lipids during the elongation step can become rate-limiting, producing yet another scaling regime as we discuss in Dear et al. (44). The *t*_1*/*2_ data observed experimentally exhibits scaling with respect to the lipid-to-protein ratio, *r*, consistent with our theoretical predictions (Fig. 4a). From these measured scaling exponents and using the theoretical scaling relationships summarized in Fig. 4b, we can reveal the reaction orders. In particular, we find that the primary nucleation of oligomers has a weak dependence on bound protein monomers with *n*_1_ ≈ 0 and a stronger dependence on free protein monomers for the oligomer nucleation with *n*_2_ ≈ 1.5. The weak dependence of primary nucleation on lipid concentration, described by *n*_1_ ≈ 0, is likely due to a saturated contribution of bound monomers to nucleation. Saturation arises because of the high relative abundance of bound monomers in the system, such that their availability is not rate-limiting. At lower lipid concentrations (*r <* 2), the decreasing availability of bound monomers decreases the primary nucleation and elongation rates and, in this regime, we find *n*_1_ *>* 0 (see Supplementary Material Sec. S4).

**Fig. 4.**
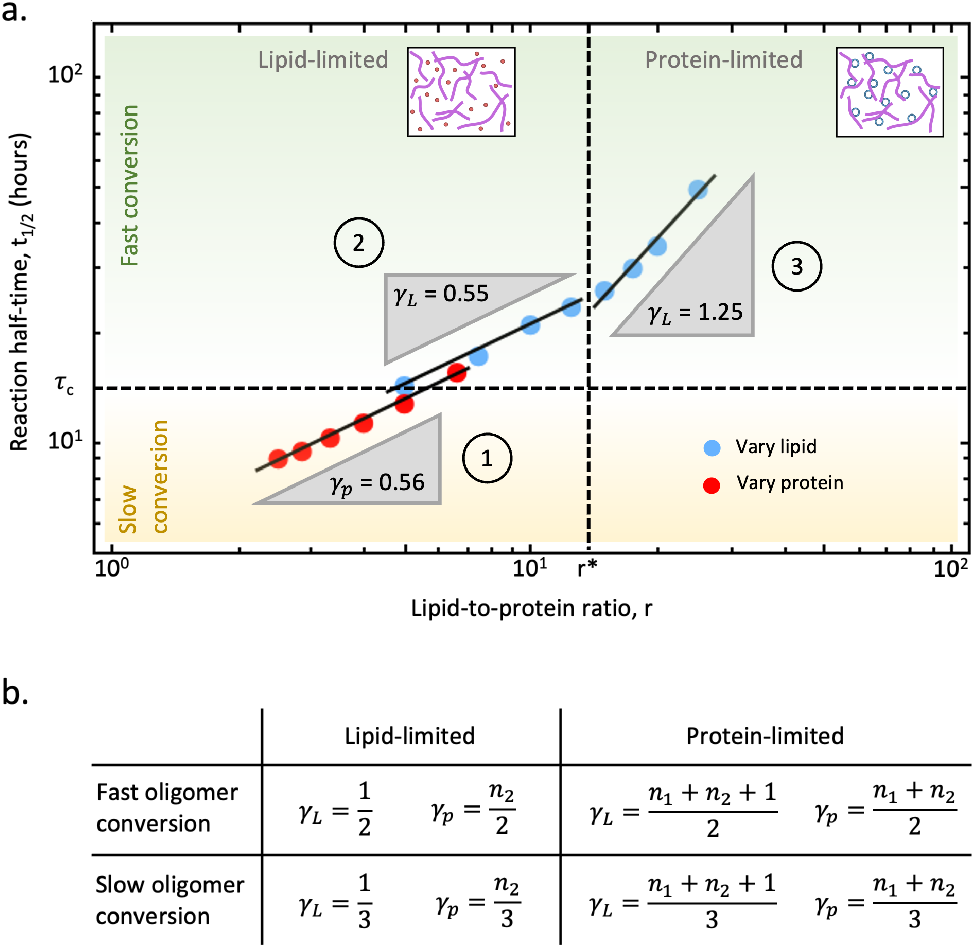
The multiphasic scaling behavior of the reaction half-time, *t*_1*/*2_, is dependent on the lipid-to-protein ratio *r* and timescale of oligomer conversion, *τ*_*c*_. **(a)** The gradient of a log-log plot of *t*_1*/*2_ against *r* reveals four distinct scaling regimes, contingent upon whether the aggregation reaction is limited by lipid or protein concentration and whether the oligomer to fibril conversion process is rapid or slow. The oligomer conversion process was determined to be rapid or slow by comparing the measured *t*_1*/*2_ with the predicted timescale of oligomer conversion, *τ*_*c*_ = ln(2)*/k*_*c*_. In region 1, the reaction is lipid-limited with a slow oligomer conversion process. The scaling exponent *γ*_*p*_ = 0.56 indicates that the nucleation of oligomers depends on free protein monomers with a reaction order *n*_2_ ≈ 1.5 (see Table in panel (b)). In region 2, the reaction is lipid-limited but the oligomer conversion process is fast. The measured scaling exponent *γ*_*L*_ = 0.55 agrees with the theoretically predicted scaling of *γ*_*L*_ = 0.5. In region 3, the reaction is protein-limited with a fast oligomer conversion process. The scaling exponent *γ*_*L*_ = 1.25 indicates that the nucleation of oligomers has a weak dependence on bound protein monomers with a reaction order *n*_1_ ≈ 0. **(b)** Summary of theoretical predictions for the scaling exponents *γ*_*p*_ and *γ*_*L*_ depending on *r* and *τ*_*c*_.

### Inhibition of α-synuclein aggregation by squalamine

Finally, we assessed the capability of our model to describe the inhibition of lipid-induced *α*-synuclein aggregation by small molecules, focusing in particular on the effects of squalamine. Squalamine, a naturally occurring aminosterol identified in the dogfish shark, inhibits lipid-induced *α*-synuclein aggregation by, as proposed previously, binding to lipids and displacing monomeric alpha-synuclein from lipid surfaces (47, 54). Fig. 5 demonstrates the impact of varying concentrations of squalamine on aggregation kinetics. Using our kinetic model we can show that the impact of squalamine on aggregation is phenomenologically consistent with an inhibition of the initial oligomer formation without explicitly modelling the effect of the inhibitor at a microscopic level (Fig. 5a). This approach allows us to determine how the associated rate constant of oligomer formation is affected by the inhibitor (Fig. 5b). This inhibition mechanism is consistent with early-time curvature changes in the kinetic profiles (Fig. 5c). Specifically, we observe that increasing the concentration of the inhibitor shifts the scaling of the aggregate mass with time from a cubic (*t*^3^) to a quadratic (*t*^2^) form. A *t*^3^ scaling is typically indicative of a two-step nucleation process with a rate-limiting oligomer conversion step, while a *t*^2^ scaling implies that the oligomer conversion process is no longer rate limiting (55). This shift indicates that the presence of squalamine slows down the overall aggregation rate by inhibiting oligomer formation, making the conversion step less rate limiting and shifting from *t*^3^ to *t*^2^ scaling. Furthermore, for squalamine to inhibit oligomer formation implies the direct involvement of a lipid surface in oligomer formation, as would be expected if oligomers were membrane-bound species. These findings are consistent with the findings that *α*-synuclein oligomers are displaced from lipid membranes by trodusquemine (56), while aminosterols like squalamine are likely capable of achieving this similar displacement effect by embedding into lipid membranes and exposing the polyamine tail (57).

**Fig. 5.**
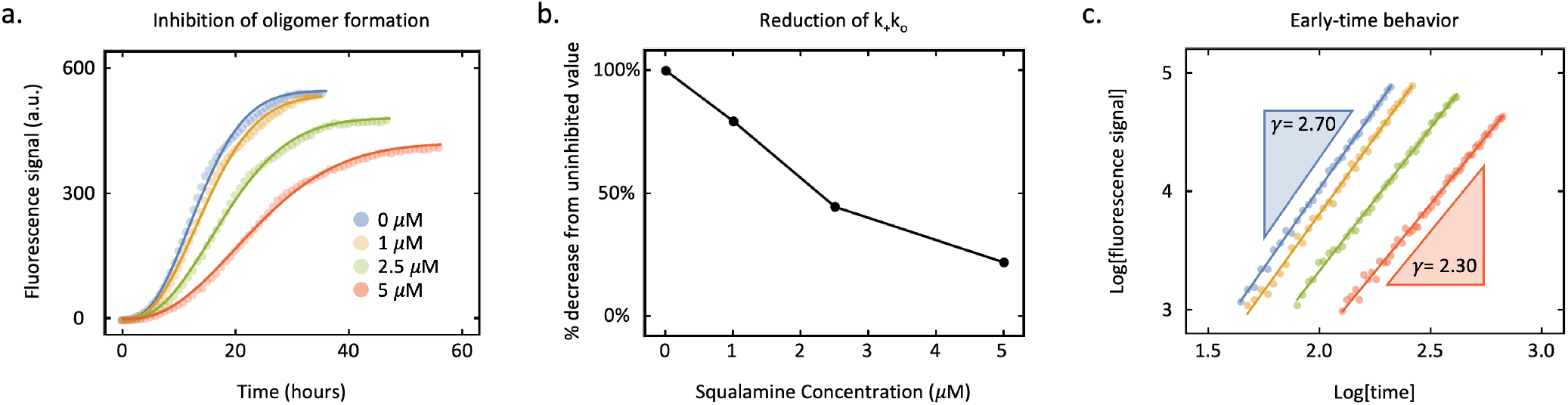
The mechanism of inhibition for lipid-induced *α*-synuclein aggregation by the small-molecule inhibitor squalamine. **(a)** Kinetics of fibril mass formation using ThT-fluorescence with constant *m*_tot_ = 100 *μ*M, *L*_tot_ = 100 *μM* and varying concentrations of squalamine (blue - 0 *μ*M, yellow - 1 *μ*M, green - 2.5 *μ*M, red - 5 *μ*M. Experimental data from Perni et al. (47). **(b)** The relative reduction in the *k*_+_*k*_*o*_ rate parameter due to increasing squalamine concentration determined by fitting analysis of the theoretical model to the experimental data in panel (a). **(c)** Early-time scaling behavior of the aggregate mass concentration with time in a double-logarithmic plot reveals a transition from *M* (*t*) ∼ *t*^3^ scaling to *M* (*t*) ∼ *t*^2^ scaling due to the formation of oligomers becoming rate-limiting, with respect to the overall two-step process, as squalamine concentration is increased.

## Conclusions

We have developed a generalizable theoretical framework to elucidate the microscopic processes that govern the dynamics of lipid-induced amyloid aggregation and derived analytical expressions for the dynamics of fibril formation. Utilizing this framework to analyze *in vitro* kinetic traces of *α*-synuclein aggregation in the presence of DMPS lipids, we have identified that lipid-induced *α*-synuclein aggregation includes two critical pathways: (1) a two-step primary nucleation process that initially forms oligomers that then convert to mature fibrils and (2) a fibril elongation process, both processes involving protein and lipid molecules and producing lipid-protein coaggregates. Our model also shows that fibril elongation has a stronger dependence on lipid concentration than primary nucleation, likely due to the lipid surface being saturated with monomer at the initial stage of the reaction when nucleation is dominant. Furthermore, our model provides essential insights into the biphasic thermodynamic behavior of the aggregation process dependent on the lipid-to-protein ratio, *r*, in particular explaining why high lipid concentrations inhibit the rate of aggregation which is not described by previous lipid-induced aggregation models (44). At low lipid-to-protein ratios, the yield of the aggregation reaction is lipid-limited, leading to a yield dependent on the initial lipid concentration. Conversely, at high lipid-to-protein ratios considered in this investigation, the fibril yield depends only on the initial protein concentration as the reaction is now protein-limited. In this regime, we found that fibril yield does not reduce as lipid concentration is raised, requiring an increasing fraction of the surface to be unbound by either monomeric or fibrillar protein at equilibrium. Lipid-protein coaggregate formation must therefore be sufficiently thermodynamically favourable to pay this energy penalty. Furthermore, the observed deceleration of the aggregation reaction at high lipid-to-protein ratios is revealed by our theoretical framework to be a natural consequence of the dilution of protein monomer surface coverage, reducing the reaction flux of fibril elongation. This framework can be applied to study other protein aggregation processes occurring in the presence of lipids, such as the aggregation of islet amyloid polypeptide and amyloid-*β*, which have been shown to interact with lipids (58, 59). Finally, our kinetic model explains existing inhibition data revealing which mechanistic steps in the aggregation reaction pathway are inhibited and quantifies the reduction in the corresponding rate parameters. These results are likely to provide essential insights into the rational design of therapeutic strategies against aberrant lipid-induced aggregation (46, 47, 60–62).

## Acknowledgments

This work was supported by ETH Zurich (AS, TCTM), the Swiss National Science Foundation (Grant Number SNS 219703 to TCTM), the Novo Nordisk Foundation (Grant Number NNF17SA0028392 to AKB and NNF20OC0059417 to CG), the Lundbeck Foundation (Grant Number R314-2018-3493 to CG) and UKRI (Grant Number 10059436 and 10061100 to MV).

## Methods

### Recombinant *α*-synuclein expression

*Escherichia coli* BL21 Gold (DE3) cells were transformed with a human *α*-synuclein-encoding pT7-7 plasmid and grown in LB (2 *× γ*T) media in the presence of ampicillin (100 *μ*g/ml). Cells were induced with 1 mM isopropyl *β*-d-1-thiogalactopyranoside (IPTG), cultured at 37°*C* overnight and harvested by centrifugation in a Beckman Avanti J-20 centrifuge with a JLA-8.1000 rotor for 20 minutes at 4000 rpm (3935*×* g) (Beckman Coulter, Fullerton, CA) and 4°*C*. The cell pellet was resuspended in 10 mM Tris–HCl (pH 7.7), 1 mM EDTA and lysed by sonication. The cell suspension was centrifuged for 20 minutes with a JA-25.50 rotor at 18 000 rpm (40000 *×* g) and 4°*C* and the supernatant was subsequently boiled by suspension in a water bath at 85 *−* 90°*C* for 20-25 minutes. The boiled supernatant was once again centrifuged for 20 minutes at 18 000 rpm (40000 *×* g) at 4°*C* to pellet heat-denatured proteins. 10 mg/mL streptomycin sulphate was added to the supernatant to precipitate DNA and rolled for 15 minutes at 4°*C* on a benchtop rolling system. To pellet precipitated DNA, the mixture was centrifuged for 20 minutes at 18 000 rpm (40000 *× g*) and 4°*C* and the supernatant was collected. To precipitate *α*-synuclein, ammonium sulphate was added to the supernatant to yield a final concentration of 361 mg/mL and the mixture was rolled for 30 minutes and 4°*C* on a benchtop rolling system before being centrifuged for 20 minutes at 18 000 rpm (40000 *× g*) at 4°*C*. The *α*-synuclein-containing pellet was resuspended in 25*mM* Tris–HCl (pH 7.7) and dialysed using a 3500 MWCO membrane in 4 L 25 mM Tris–HCl (pH 7.7). *α*-synuclein was purified by ion exchange on a Q-Sepharose HP HiScaleTM 26/20 column (Cytiva, USA) before size exclusion chromatography (SEC) on a HiLoadTM 16/600 Superdex 75 pg column (Cytiva, USA) into the appropriate experimental buffer. To determine the concentrations in solution, the absorbance value of the protein was measured at 275 nm and an extinction coefficient of 5600 M^−1^. The protein solutions were divided into aliquots, flash frozen in liquid nitrogen and stored at *−* 80°*C* until required for use. Before experimental use, the protein was thawed and size excluded again on a Superdex 75 10/300 GL column (Cytiva, USA) into the appropriate experimental buffer to remove any aggregate species that may have formed during freezing, after which the concentration of the monomeric elution was measured on a NanoDrop One (Thermo Scientific, USA) at 260*/*280 nm and again with an extinction coefficient of 5600 M^−1^.

### Lipid-induced aggregation assay

#### Lipid vesicle preparation

Powder of 1,2-dimyristoyl-sn-glycero-3-phospho-L-serine (DMPS) was dissolved to a final concentration of 5*μ*M in 20 mM sodium phosphate buffer at pH 6.5 and stirred at 900 rpm at 50°*C* for 2-2.5 hours followed by five freeze-thaw cycles in dry ice and a hot water bath at 45°*C* to ensure unilamerallity. To form vesicles, the solution was sonicated by a Sonopuls HD 2070 probe sonicator (Bandelin) for 3 × 5 minute cycles on a 50% cycle at 10% maximum power, and centrifuged at 15 000 rpm for 30 minutes at 25°*C* to remove residue formed during sonication.

#### Sample preparation and measurement of α-synuclein lipid-induced aggregation kinetics

Monomeric *α*-synuclein was size excluded into 20 mM sodium phosphate buffer at pH 6.5 and the concentration was measured. The protein was diluted to the desired concentration in 20 mM sodium phosphate buffer at pH 6.5, and combined with 50 *μ*M Thioflavin-T (ThT). DMPS vesicles were added to induce the aggregation to obtain a final concentration of 100 *μ*M vesicles. The change in the ThT fluorescence signal was monitored at 30°*C* using a Fluostar

Optima or Polarstar Omega fluorescence plate reader (BMG Labtech, Aylesbury, UK) in bottom reading mode under quiescent conditions over the time course indicated with an excitation filter at 440 nm and an emission filter at 480 nm. Corning 96-well plates with half-area (3881, polystyrene, black with clear bottom) non-binding surfaces sealed with aluminium sealing tape were used for each experiment.

#### Kinetic equations for lipid-induced aggregation

Here, we provide the mathematical details of our kinetic model describing lipid-induced aggregation as a surface reaction. In Supplementary Material Sec. S3, we provide an introduction to chemical kinetics on surfaces. Here, we adopt the same tools for lipid-induced aggregation and describe the process in terms of differential equations for the surface coverages of monomers and aggregates. These describe the fractions of lipid surface that are occupied by monomers or aggregates, respectively, and are defined as

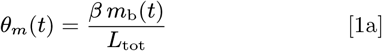

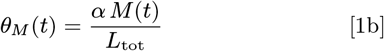

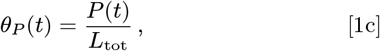

where *m*_b_ is the concentration of monomers bound to the surface, *P*(*t*) and *M*(*t*) denote the number and mass concentrations of fibrils, *L*_tot_ is the total concentration of lipids, *α* is the average number of lipid units bound by a single protein monomer and *β* is the average number of lipid units bound by a single protein monomer in fibril form. Supplementary Material Table S2 summarizes the definitions of these various parameters used in our theoretical models of lipid-induced protein aggregation.

#### Kinetic equations for one-step primary nucleation model

In the simplest setting, fibrils are formed via one-step primary nucleation and grow by lipid-induced elongation. This scenario is described by the following kinetic equations

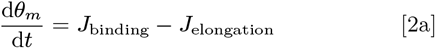

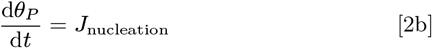

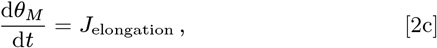

where *J* denotes the reaction flux for the different processes, including monomer binding/unbinding from lipids, fibril nucleation and elongation. The explicit expression for the monomer binding flux *J*_binding_ is:

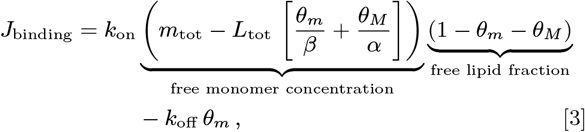

when *k*_on_ is the rate constant of protein monomer binding to the lipid surface and *k*_off_ is the rate constant of protein monomer dissociation from the lipid surface. The expression for the lipid-induced elongation flux is:

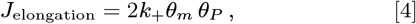

where *k*_+_ is the rate constant of elongation. For the flux of one-step primary nucleation, we have:

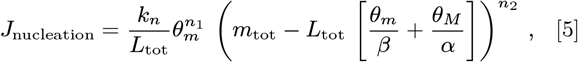

where *k*_*n*_ is the rate constant of primary nucleation of fibrils. The reaction orders *n*_1_ and *n*_2_ allow for different possible contributions of free and bound monomers to the primary nucleation rate, respectively.

#### Kinetic equations for two-step primary nucleation model

To extend Eq. Eq. (2) to account for a two-step primary nucleation mechanism, we consider the concentration of intermediate oligomers that then convert to mature, elongation-capable, fibrils. The following set of differential equations then describes the time evolution of oligomer concentration and the surface coverage terms:

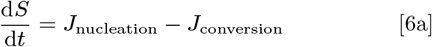

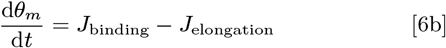

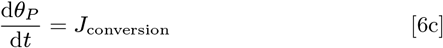

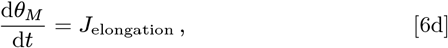

where *S* is the concentration of oligomers and *J*_conversion_ is the reaction flux describing the conversion of oligomers to fibrils. Similar to the one-step primary nucleation flux, we have:

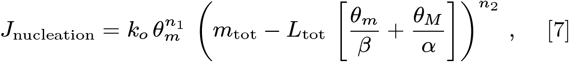

where *k*_*o*_ is now the rate constant of primary nucleation of oligomers. The oligomer conversion flux is:

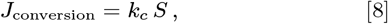

where *k*_*c*_ is the rate constant of oligomer conversion.

[8]

#### Self-consistent solutions to aggregation kinetics

We used self-consistent methods and asymptotic approaches (40) to derive analytical solutions to the kinetic Eqs. Eq. (2) and Eq. (6). In Supplementary Material Sec. S3, we discuss these techniques in detail in the context of surface reactions and here we apply these tools to Eqs. Eq. (2) and Eq. (6). The first step is to consider the limit when there is a timescale separation between the rapid binding dynamics of proteins to the lipid surface and slower aggregation kinetics. In this limit, we can assume a pre-equilibrium between free and bound monomers at all times. In particular, at *t* = 0, the fraction of *α*-synuclein bound to lipids is 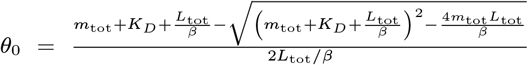 where 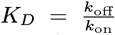 is the equilibrium dissociation constant. To transform the differential equation system into a fixed point problem, we integrate Eq. Eq. (2c) resp. Eq. (6d) such that they are of the form 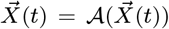 for some operator 𝒜with 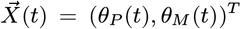 for the one-step primary nucleation as well as the secondary nucleation model and 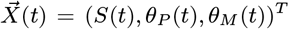 for the two-step primary nucleation model. This procedure yields the following fixed-point equation

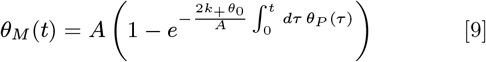

where

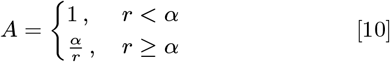

This problem is solved self-consistently if a fixed point 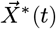 of the operator 𝒜 satisfying 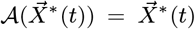 is found. According to the contract mapping principle, for an initial guess 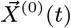 sufficiently close to the fixed point 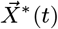, the fixed point can be computed iteratively as 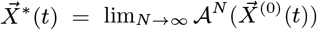. Our approach includes linearizing the kinetic equations for small times *t* and using the linearized solutions as a starting point 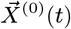 for obtaining increasingly accurate self-consistent solutions by repeatedly applying the fixed point operator 𝒜. In all cases, we find that the analytical expression for the fibril mass fraction can be written as

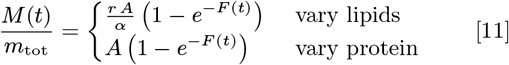

where the explicit form of *F*(*t*) depends on the dominant mechanism of fibril formation (see below for explicit expressions).

#### One-step primary nucleation

For one-step primary nucleation, the early-time expression for *θ*_*P*_ (*t*) is

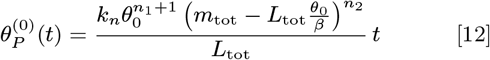

Using this solution as an initial guess in Eq. Eq. (11), we find the function *F*(*t*) after one fixed-point iteration is given by

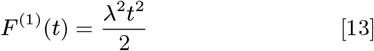

where

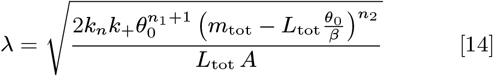

is an effective rate constant of aggregate proliferation through primary nucleation and growth. Therefore, there is only one fitting rate parameter, which is the combination *k*_+_*k*_*n*_. Applying this procedure up to third order, we obtain

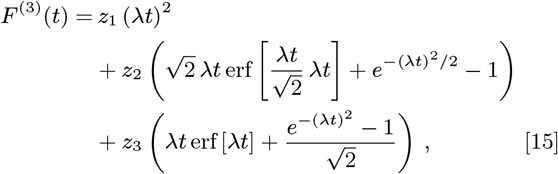

where 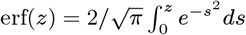 is the error function and the constants *z*_1_, *z*_2_ and *z*_*3*_are defined as

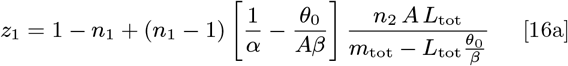

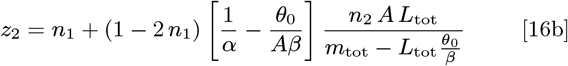

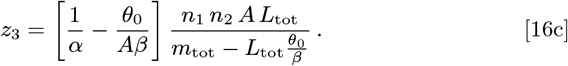

#### Two-step primary nucleation

For two-step primary nucleation, the early-time solution for *θ*_*P*_ is

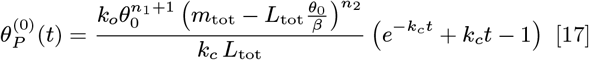

Integrating this expression, we find *F*(*t*) is given by

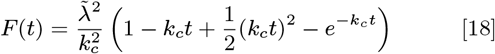

where

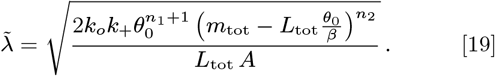

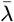 is the same as *λ* exept that *k*_*n*_ is replaced by *k*_*o*_. The fitting rate parameters are in this case *k*_+_*k*_*o*_ and *k*_*c*_.

#### Steady-state behaviour

Using Eq. Eq. (11), we find the steady-state plateau value to be

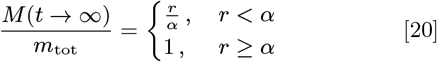

where we used *F*(*t* → ∞) = ∞. This yields the plots in Figs. 2b, c.

