## Supplementary Information for "Global kinetic model of lipid-induced *α*-synuclein aggregation and its inhibition by small molecules"

**S4 Low lipid-to-protein ratios** **23**

**S5 Cooperative binding of protein monomer to lipid surface** **24**

### S1. Supplementary Figures

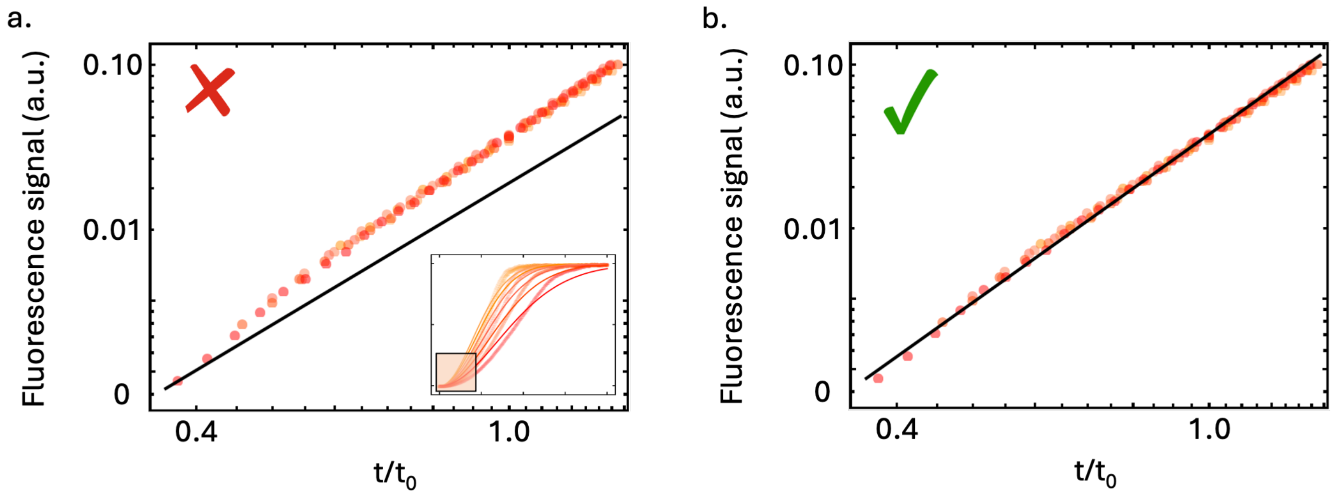

**Fig. S1.** Early time behavior (up to tenth-time $t_{10\%}$ and normalized by $t_0 = t_{5\%}$) in double-logarithmic plot shows higher than $t^2$ behaviour (solid line in (a): $M \sim t^2$, solid line in (b): $M \sim t^{2.4}$), indicating the presence of a non-negligible conversion step of oligomeric species to growth-competent fibrils in the primary nucleation process.

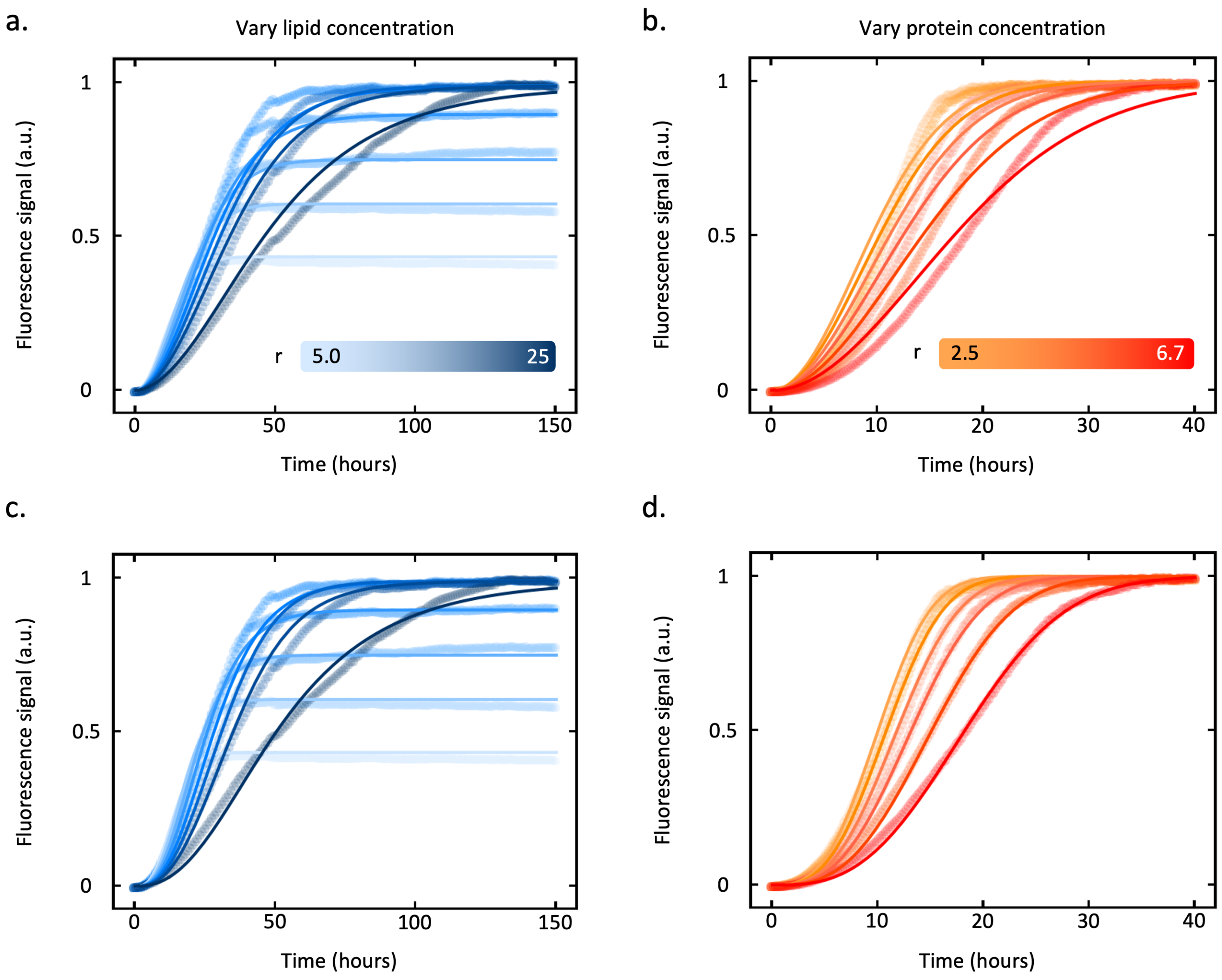

**Fig. S2.** Best global fits of experimental data shown in Fig. 3 of the main text to the **numerically solved** chemical kinetics model with: one-step primary nucleation **(a)** and **(b)** and two-step primary nucleation **(c)** and **(d)**.

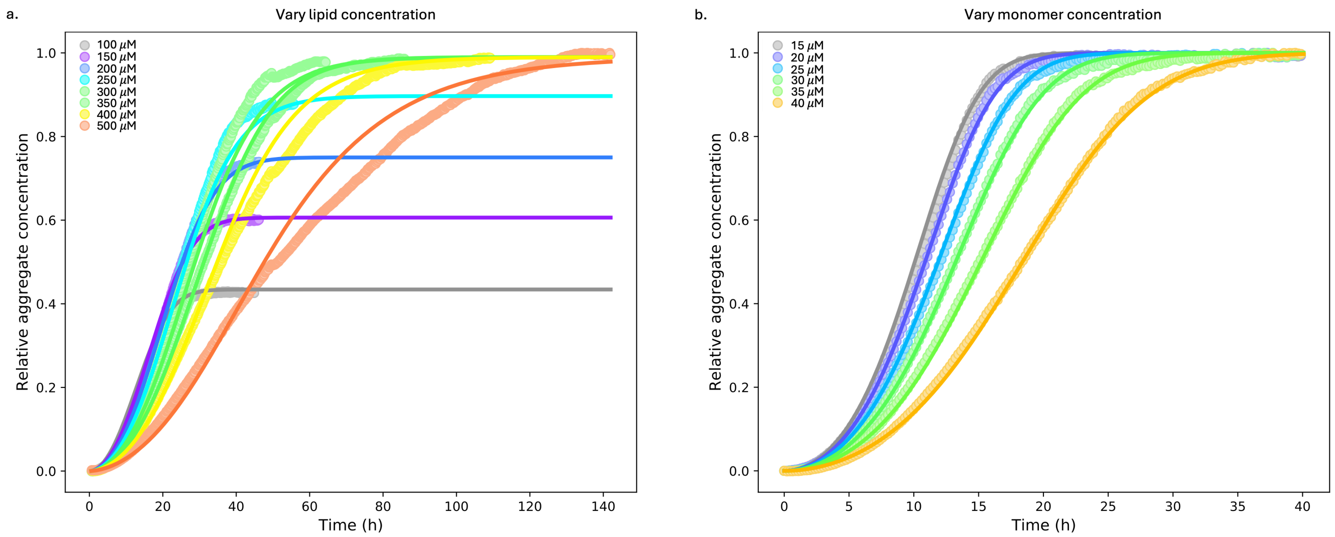

**Fig. S3.** Best global fits of experimental data shown in Fig. 3 of the main text to a model that includes secondary nucleation mechanisms.

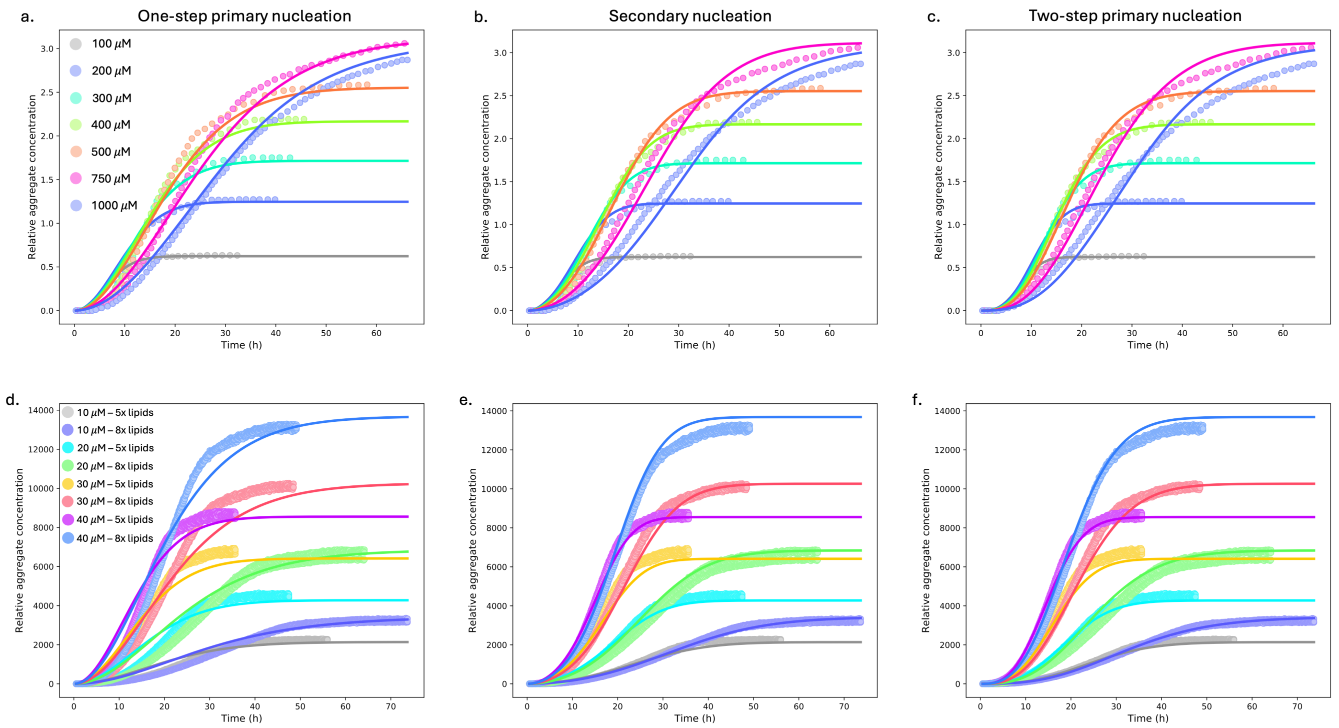

**Fig. S4.** Best global fits of previously published experimental data of $\alpha$-synuclein aggregation in the presence of DMPS SUVs, where the top panels are data from (1) and the bottom panels are data from (2). The fits were found using the one-step primary nucleation model (panels a and d), secondary nucleation model (b and e), and two-step nucleation model (panels c and f).

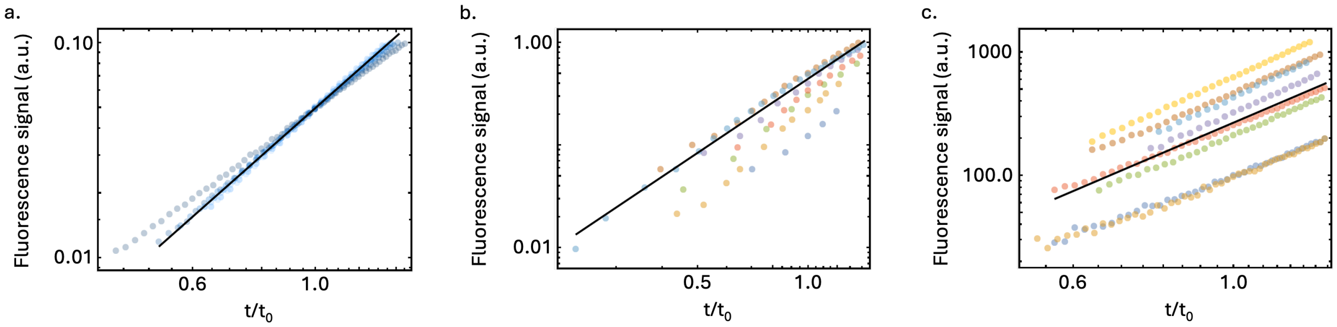

**Fig. S5.** Early time behaviour of *in vitro* lipid-induced $\alpha$-synuclein aggregation in double-logarithmic plots. **(a)** Data from Fig. 3a, c up to tenth-time $t_{10\%}$ and normalized by $t_0 = t_{5\%}$, with the solid line plotting $M \sim t^{2.275}$. **(b)** Data from (1) up to $t_{33\%}$ and normalized by $t_0 = t_{16.7\%}$, with the solid line plotting $M \sim t^{2.4}$. **(c** Data from (2) up to $t_{10\%}$ and normalized by $t_0 = t_{5\%}$, with the solid line plotting $M \sim t^{2.5}$. As all exponents are greater than 2, this indicates the presence of a conversion step in the primary nucleation of fibrils.

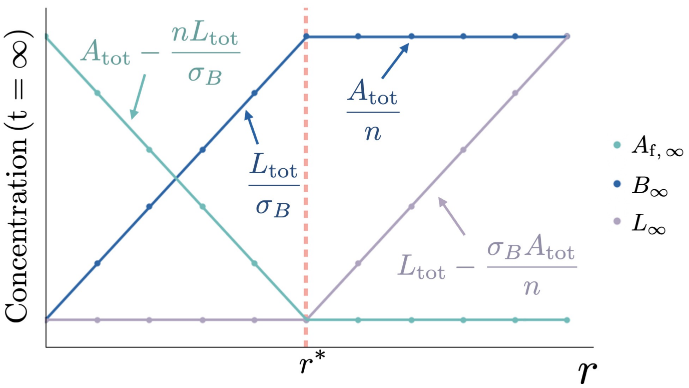

**Fig. S6.** Steady state behaviour of free reactant ($A_{\mathrm{f}}(t = \infty)$), product ($B(t = \infty)$) and free lipid site ($L(t = \infty)$) concentrations in the surface and reactant limited regimes of the system's behaviour, with the bifurcation point in behavior occurring at $r^* = \sigma_B/n$. The solid lines are the theoretical curves Eq. (S8), while the data points are obtained from numerical simulations of Eq. (S7).

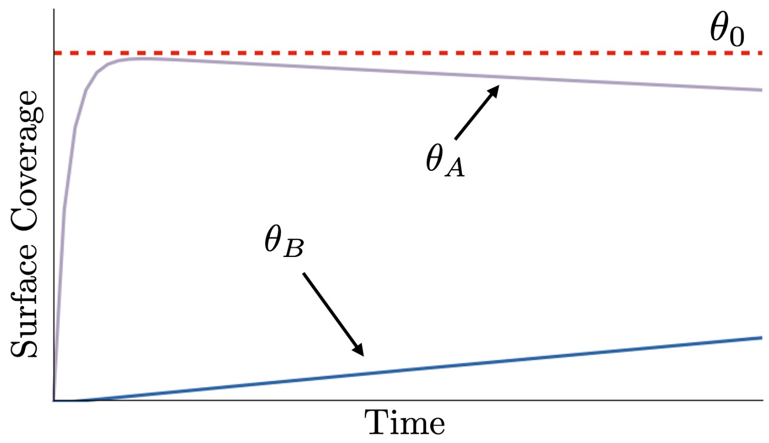

**Fig. S7.** Dynamics of surface coverage terms, $\theta_A$ and $\theta_B$, predicted by numerically evaluating Eq. (S7) compared to the approximated early-time solution found to describe a constant bound concentration/surface coverage of specie A (Eq. (S11)).

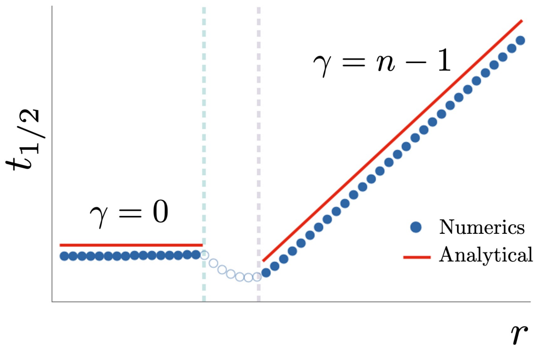

**Fig. S8.** A comparison of $t_{1/2}$ values, from analytical expressions derived in Section S3.5 and numerical evaluation of the differential equation system (Eq. (S7)). This revealed the scaling dependency on $r$ in the surface limited regime ($r < r^*$) and reactant limited regime ($r > r_s$), where the intermediate region defined as $r^* < r < r_s$ does not provide relevant insight into the parameter values under investigation. On a Log-Log plot, the gradient enables the reaction order to be quantified, $n$, using the scaling results summarized in Table S8.

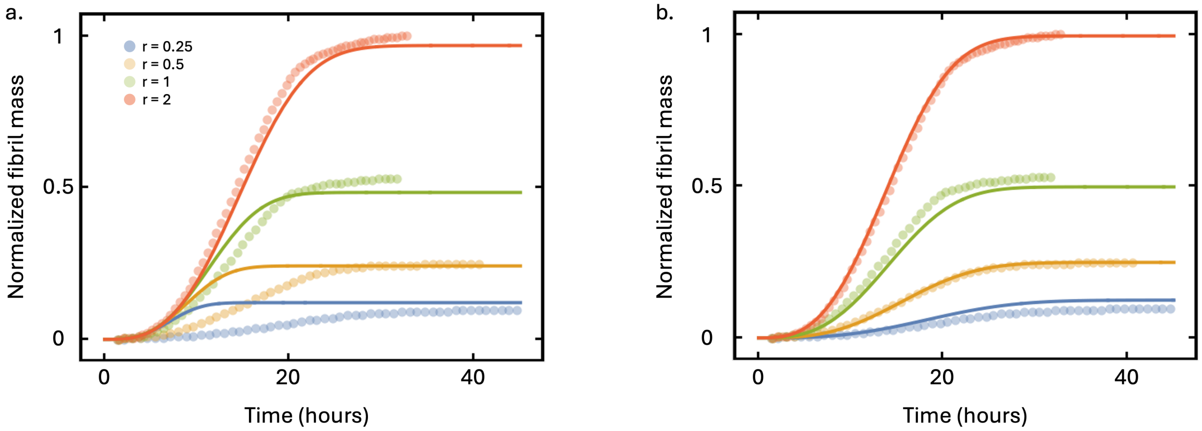

**Fig. S9.** Effect of low lipid-to-protein ratios on lipid-induced $\alpha$-synuclein aggregation kinetics. **(a)** Experimental data from (2) and solid lines showing the best global fit to the analytical solution of the two-step primary nucleation model. **(b)** Solid lines showing the best global fit to the analytical solution of the two-step primary nucleation model with the low ratio extension added (described in Supplementary Material Sec. S4).

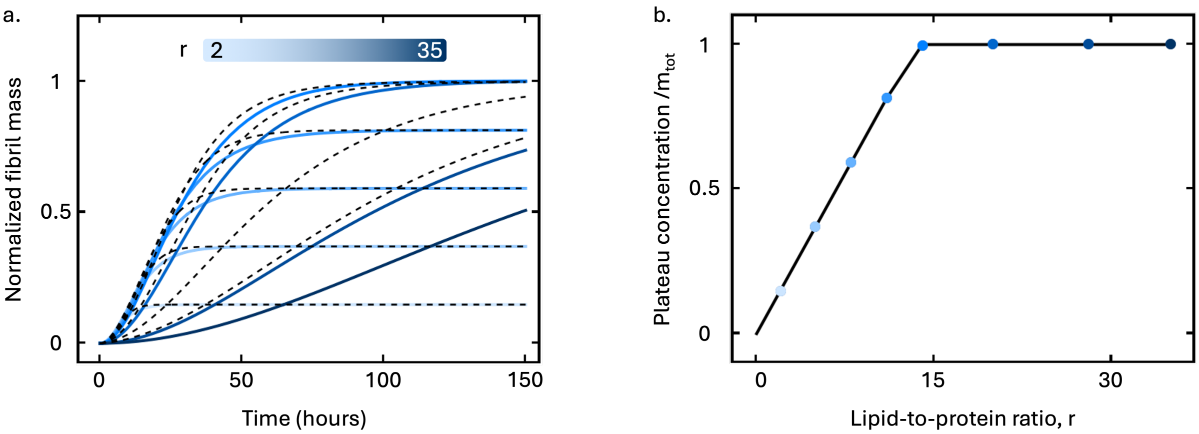

**Fig. S10.** Effect of cooperative protein binding on the kinetic and steady-state behavior of the theoretical model. **(a)** Dashed lines show the predicted kinetic traces from the theoretical model with non-cooperative protein binding to the lipid surface, solid lines are with cooperative binding ($n_{\mathrm{b}} = 3$). **(b)** Solid line shows the predicted plateau concentration of fibril mass with non-cooperative binding, and the coloured markers show the predicted plateau concentration with cooperative binding ($n_{\mathrm{b}} = 3$) having no effect of the system's steady state behavior.

### S2. Supplementary Tables

**Table S1.** Definitions of the parameters used in Section S3. We use italicized letters to denote concentrations and roman letters to denote the associated species.

| Symbol | Definition |
|---|---|
| $A_{\mathrm{tot}}$ | Total initial concentration of reactant molecules |
| $L_{\mathrm{tot}}$ | Total initial concentration of binding sites |
| $\theta_A$ | Surface coverage of reactant molecules on the surface |
| $A_{\mathrm{b}}$ | Concentration of reactant molecules bound to the surface |
| $A_{\mathrm{f}}$ | Concentration of reactant molecules not bound to the surface |
| $\theta_B$ | Surface coverage of product molecules on the surface |
| $B$ | Concentration of product molecules |
| $\sigma_A$ | Average number of surface units bound by a single reactant molecule |
| $\sigma_B$ | Average number of surface units bound by a single product molecule |
| $k_{\mathrm{on}}$ | Rate of free reactant binding to the surface |
| $k_{\mathrm{off}}$ | Rate of bound reactant dissociation from the surface |
| $K_{\mathrm{D}}$ | Dissociation constant of reactant-surface binding |
| $k_{\mathrm{n}}$ | Rate constant describing the conversion of reactant to product |
| $n$ | Reaction order describing the dependence of the reaction on bound reactant molecules |
| $\theta_0$ | Approximated solution for $\theta_A$ applicable at early-time scales |
| $t_{1/2}$ | Time for half the total concentration of reactants to convert to product form |

**Table S2.** Definitions of all symbols used in the theoretical modelling of lipid-induced $\alpha$-synuclein aggregation.

| Symbol | Definition |
|---|---|
| $m_{\mathrm{tot}}$ | Total initial concentration of protein monomers |
| $L_{\mathrm{tot}}$ | Total initial concentration of lipids |
| $\theta_m$ | Surface coverage of protein monomers on the lipid surface |
| $m_{\mathrm{b}}$ | Mass concentration of protein monomers bound to the lipid surface |
| $m_{\mathrm{f}}$ | Mass concentration of protein monomers not bound to the lipid surface |
| $\theta_P$ | Probability of meeting a fibril end on the lipid surface |
| $P$ | Number concentration of protein fibrils |
| $\theta_M$ | Surface coverage of protein monomers in fibril form on the lipid surface |
| $M$ | Mass concentration of protein monomers in fibril form |
| $\beta$ | Average number of lipid units bound by a single protein monomer |
| $\alpha$ | Average number of lipid units bound by a single protein monomer in fibril form |
| $k_{\mathrm{on}}$ | Rate of free monomer binding to the lipid surface |
| $k_{\mathrm{off}}$ | Rate of bound monomer dissociation from the lipid surface |
| $K_{\mathrm{D}}$ | Dissociation constant of protein-lipid binding |
| $k_+$ | Rate constant describing protein fibril elongation |
| $k_n$ | Rate constant describing primary nucleation of protein fibrils |
| $k_o$ | Rate constant describing primary nucleation of protein oligomers |
| $k_c$ | Rate constant describing the conversion of protein oligomers to fibrils |
| $k_2$ | Rate constant describing secondary nucleation of protein fibrils |
| $n_1$ | Reaction order parameter describing the dependence of primary nucleation on bound monomers |
| $n_2$ | Reaction order parameter describing the dependence of primary nucleation on free monomers |
| $n_3$ | Reaction order parameter describing the dependence of secondary nucleation on bound monomers |
| $n_4$ | Reaction order parameter describing the dependence of secondary nucleation on free monomers |
| $\theta_0$ | Approximated solution for $\theta_m$ applicable at early-time scales |
| $t_{1/2}$ | Time for half the total concentration of protein monomers to convert to fibril form |
| $\tau_c$ | Timescale for conversion of oligomers to fibrils |

**Table S3.** Summary of fitting parameters used to describe the experimental data with the **analytical solution** to the chemical kinetics model with one-step primary nucleation in Figs. 3a and 3b of the main text.

| Parameter | Vary lipid concentration (Fig. 3a) | Vary monomer concentration (Fig. 3b) |
|---|---|---|
| $\sqrt{k_+ k_n}$ | 0.16 $\text{hr}^{-1}$ | 0.14 $\text{hr}^{-1}$ |
| $\alpha$ | 13.6 (average) | 11.4 |
| $n_1$ | 0 | 0 |
| $n_2$ | 1 | 1 |

**Table S4.** Summary of fitting parameters used to describe the experimental data with the **numerically solved** chemical kinetics model with one-step primary nucleation in Supplementary Material Figs. S2a and S2b.

| Parameter | Vary lipid concentration (Fig. 3a) | Vary monomer concentration (Fig. 3b) |
|---|---|---|
| $k_+$ | 1 $\text{hr}^{-1}$ | 1 $\text{hr}^{-1}$ |
| $k_{on}$ | 0.25 $M^{-1}$ $\text{hr}^{-1}$ | 0.25 $M^{-1}$ $\text{hr}^{-1}$ |
| $\sqrt{k_+ k_n}$ | 0.18 $\text{hr}^{-1}$ | 0.15 $\text{hr}^{-1}$ |
| $\alpha$ | 13.6 (average) | 11.4 |
| $n_1$ | 0 | 0 |
| $n_2$ | 1 | 1 |

**Table S5.** Summary of fitting parameters used to describe the experimental data with the **analytical solution** to the chemical kinetics model with two-step primary nucleation in Figs. 3c and 3d of the main text.

| Parameter | Vary lipid concentration (Fig. 3c) | Vary monomer concentration (Fig. 3d) |
|---|---|---|
| $\sqrt{k_+ k_o}$ | 0.11 $\text{hr}^{-1}$ | 0.16 $\text{hr}^{-1}$ |
| $k_c$ | 0.26 $\text{hr}^{-1}$ | 0.042 $\text{hr}^{-1}$ |
| $\alpha$ | 13.6 (average) | 11.4 |
| $n_1$ | 0 | 0 |
| $n_2$ | 1.5 | 1.5 |

**Table S6.** Summary of fitting parameters used to describe the experimental data with the **numerically solved** chemical kinetics model with two-step primary nucleation in Supplementary Material Figs. S2c and S2d.

| Parameter | Vary lipid concentration (Fig. 3c) | Vary monomer concentration (Fig. 3d) |
|---|---|---|
| $k_+$ | 1 $\text{hr}^{-1}$ | 1 $\text{hr}^{-1}$ |
| $k_{on}$ | 0.25 $M^{-1}$ $\text{hr}^{-1}$ | 0.25 $M^{-1}$ $\text{hr}^{-1}$ |
| $\sqrt{k_+ k_o}$ | 0.13 $\text{hr}^{-1}$ | 0.15 $\text{hr}^{-1}$ |
| $k_c$ | 0.11 $\text{hr}^{-1}$ | 0.051 $\text{hr}^{-1}$ |
| $\alpha$ | 13.6 (average) | 11.4 |
| $n_1$ | 0 | 0 |
| $n_2$ | 1.5 | 1.5 |

**Table S7.** Summary of fitting parameters used to describe the experimental data with the **numerically solved** chemical kinetics model with secondary nucleation in Supplementary Material Fig. S3.

| Parameter | Vary lipid concentration (Fig. S2a) | Vary monomer concentration (Fig. S2b) |
|---|---|---|
| $k_+$ | 1 $\mathrm{hr}^{-1}$ | 1 $\mathrm{hr}^{-1}$ |
| $k_{on}$ | 0.25 $M^{-1}$ $\mathrm{hr}^{-1}$ | 0.25 $M^{-1}$ $\mathrm{hr}^{-1}$ |
| $\sqrt{k_+ k_n}$ | 0.14 $\mathrm{hr}^{-1}$ | 0.073 $\mathrm{hr}^{-1}$ |
| $\sqrt{k_+ k_2}$ | 0.11 $\mathrm{hr}^{-1}$ | 0.11 $\mathrm{hr}^{-1}$ |
| $\alpha$ | 13.6 (average) | 11.4 |
| $n_1$ | 0 | 0 |
| $n_2$ | 1 | 1 |
| $n_3$ | 0 | 0 |
| $n_4$ | 1 | 1 |

**Table S8.** Summary of $\alpha$ values used to fit the vary lipid concentration data (Figs. 3a, 3c and Supplementary Material Figs. S2a, S2c and S3a) with the different kinetic models ($\alpha = 11.4$ was used to fit all vary protein concentration data). This range of values is reasonable based on previous analysis suggesting $10 < \alpha < 15$ depending on lipid concentration (2).

| $L_{\mathrm{tot}}$ | $\alpha$ |
|---|---|
| 100 $\mu M$ | 11.4 |
| 150 $\mu M$ | 12.25 |
| 200 $\mu M$ | 13.2 |
| 250 $\mu M$ | 13.8 |
| 300 $\mu M$ | 14.5 |
| 350 $\mu M$ | 14.5 |
| 400 $\mu M$ | 14.5 |
| 500 $\mu M$ | 14.5 |

**Table S9.** Summary of fitting parameters used to describe the experimental data of aggregation inhibition by squalamine with the **analytical solution** to the chemical kinetics model with two-step primary nucleation in Fig. 5a.

| [squalamine] | $\sqrt{k_+ k_o}$ | $k_c$ | $\alpha$ | $n_1$ | $n_2$ | $L_{\text{tot}}$ |
|---|---|---|---|---|---|---|
| $0\,\mu M$ | 0.025 hr$^{-1}$ | 0.41 hr$^{-1}$ | 10.5 | 0 | 1.5 | $100\,\mu M$ |
| $1\,\mu M$ | 0.022 hr$^{-1}$ | 0.41 hr$^{-1}$ | 10.5 | 0 | 1.5 | $98\,\mu M$ |
| $2.5\,\mu M$ | 0.016 hr$^{-1}$ | 0.41 hr$^{-1}$ | 10.5 | 0 | 1.5 | $88\,\mu M$ |
| $5\,\mu M$ | 0.012 hr$^{-1}$ | 0.41 hr$^{-1}$ | 10.5 | 0 | 1.5 | $77\,\mu M$ |

**Table S10.** Summary of fitting parameters used to describe the experimental data with the **analytical solution** to the chemical kinetics model with two-step primary nucleation in Supplementary Material Fig. S9, where the analytical solution has been modified to examine low lipid-to-protein ratios (see Supplementary Material Sec. S4).

| Parameter | Vary lipid concentration misfit (Supplementary Material Fig. S9a) | Vary lipid concentration (Supplementary Material Fig. S9b) |
|---|---|---|
| $\sqrt{k_+ k_o}$ | 0.18 $\text{hr}^{-1}$ | 0.21 $\text{hr}^{-1}$ |
| $k_c$ | 0.0045 $\text{hr}^{-1}$ | 0.016 $\text{hr}^{-1}$ |
| $\alpha$ | 10.5 | 10.5 |
| $n_1$ | 1 | 1 |
| $n_2$ | 1.5 | 1.5 |

### S3. Chemical kinetics on surfaces: A review

In this section, we review some fundamental concepts and tools from the theory of chemical kinetics on surfaces (3). This section summarises the key results that are used in the analysis of protein aggregation on lipid surfaces and provides an introduction for readers who are not familiar with these concepts. As an illustrative example, we will consider the chemical reaction $nA \rightarrow B$, where the reactant $A$ is able to bind to and dissociate from a surface and the product $B$ remains bound and can be described with the following multi-step reaction pathway:

$$\mathrm{A_f} + \mathrm{L} \underset{k_{\mathrm{off}}}{\overset{k_{\mathrm{on}}}{\rightleftharpoons}} \mathrm{nA_b} \xrightarrow{\mathrm{k}_n} \mathrm{B},$$

where $\mathrm{A_f}$ and $\mathrm{A_b}$ are the free and bound reactant respectively, B is the product molecule and L is the free binding sites on the surface. The rate constants, $k_{\mathrm{on}}$ and $k_{\mathrm{off}}$, are the rates of binding and dissociation of A to and from the surface respectively. The rate of conversion of A to B is $k_{\mathrm{n}}$ and the reaction order is $n$, and we assume that the product B stays bound to the surface without unbinding.

**S3.1. Surface coverage vs bulk concentration.** The key quantities needed to describe the kinetics of chemical reactions on surfaces are the surface coverages of reactants and products. These are defined by

$$\theta_A = \frac{\sigma_A A_{\mathrm{b}}}{L_{\mathrm{tot}}} \quad \text{[S1a]}$$

$$\theta_B = \frac{\sigma_B B}{L_{\mathrm{tot}}}\,, \quad \text{[S1b]}$$

where $\theta_A$ and $\theta_B$ are the non-dimensional surface coverage of A and B respectively, taking values between 0 and 1. $L_{\mathrm{tot}}$ is the total concentration of surface sites. $\sigma_A$ and $\sigma_B$ represent the number of sites bound to a molecule $A$ and $B$ respectively. Note that the fraction of free sites is:

$$\theta_L = 1 - \theta_A - \theta_B\,, \quad \text{[S2]}$$

which follows from the conservation of surface coverage terms in the system $\theta_A + \theta_B + \theta_L = 1$.

**S3.2. Kinetic equations on surfaces.** To describe the kinetics of a chemical reaction $n\mathrm{A} \rightarrow \mathrm{B}$, we first describe the binding-dissociation kinetics of A in terms of bulk concentrations $A_{\mathrm{b}}$, $A_{\mathrm{f}}$ and $L$:

$$\left.\frac{\mathrm{dA_b}}{\mathrm{dt}}\right|_{\mathrm{binding}} = k_{\mathrm{on}} A_{\mathrm{f}} \frac{L}{\sigma_A} - k_{\mathrm{off}} A_{\mathrm{b}}\,, \quad \text{[S3]}$$

where these quantities can be described in terms of the surface coverage through the following relationships:

$$L = L_{\mathrm{tot}}(1 - \theta_A - \theta_B) \quad \text{[S4a]}$$

$$A_{\mathrm{f}} = A_{\mathrm{tot}} - A_{\mathrm{b}} - nB\,, \quad \text{[S4b]}$$

where $A_{\mathrm{tot}}$ is the total concentration of A. Using this knowledge, Eq. (S3) can be modified to describe the kinetics of $\theta_A$ as:

$$\frac{L_{\mathrm{tot}}}{\sigma_A}\frac{\mathrm{d}\theta_A}{\mathrm{dt}} = k_{\mathrm{on}}\left(A_{\mathrm{tot}} - L_{\mathrm{tot}}\left[\frac{\theta_A}{\sigma_A} + \frac{n\theta_B}{\sigma_B}\right]\right)(1 - \theta_A - \theta_B)\frac{L_{\mathrm{tot}}}{\sigma_A} - k_{\mathrm{off}}\frac{L_{\mathrm{tot}}}{\sigma_A}\theta_A\,, \quad \text{[S5]}$$

where terms are multiplied by $L_{\mathrm{tot}}/\sigma_A$ to convert from concentration to surface coverage. We assume that as the conversion of A to B occurs on the surface, the rate of this reaction is proportional to the surface coverage of A, rather than the concentration. This is physically interpretable as the reaction occurs when bound A molecules meet on the surface, which is dependent on the coverage of A on that surface. The reaction flux of the conversion of $A$ to $B$ can then be described as:

$$\left.\frac{\mathrm{d}\theta_{\mathrm{A}}}{\mathrm{dt}}\right|_{\mathrm{conversion}} = nk_n\theta_A^n\,. \quad \text{[S6]}$$

Combining Eq. (S5) and Eq. (S6) yields the differential equation system now describing surface coverage dynamics:

$$\frac{\mathrm{d}\theta_{\mathrm{A}}}{\mathrm{dt}} = \underbrace{k_{\mathrm{on}}\left(A_{\mathrm{tot}} - L_{\mathrm{tot}}\left[\frac{\theta_A}{\sigma_A} + \frac{n\theta_B}{\sigma_B}\right]\right)(1 - \theta_A - \theta_B) - k_{\mathrm{off}}\theta_A}_{\text{Binding/dissociation kinetics}} - \underbrace{nk_{\mathrm{n}}\theta_A^n}_{\text{Reaction}} \quad \text{[S7a]}$$

$$\frac{\mathrm{d}\theta_{\mathrm{B}}}{\mathrm{dt}} = k_{\mathrm{n}}\theta_A^n \quad \text{[S7b]}$$

**S3.3. Steady-state analysis.** The steady-state behaviour of this system is determined either by the conversion of A to B stopping due to the consumption of all available surface sites (surface limited regime), or the surface sites are available in excess and all A is converted to B (reactant limited regime). To understand the origin of this biphasic behaviour, we examined the steady states of the kinetic equations. These were obtained by setting the time derivatives to zero in Eq. (S7). At the end of the reaction $\theta_A(t = \infty) = 0$, as any bound reactants are converted to product. From Eq. (S7a) we see that the two fixed points for the product surface coverage are at: $\theta_B(t = \infty) = \alpha\, A_{\rm tot}/n\, L_{\rm tot}$ and $\theta_B(t = \infty) = 1$. These correspond to two possible outcomes of the reaction with either $B(t = \infty) = A_{\rm tot}/n$ or $B(t = \infty) = L_{\rm tot}/\sigma_B$. Since the final product concentration $B(t = \infty)$ cannot exceed the available reactant concentration $A_{\rm tot}/n$ (normalized by reaction order), the second fixed point is stable only if $r < \sigma_B/n$, where $r = L_{\rm tot}/A_{\rm tot}$. Hence, depending on the value or $r$, the system will evolve towards a different fixed point. In summary, the values of the relevant quantities at steady state for these two fixed points are:

$$L(t=\infty) = \begin{cases} 0, & r < \sigma_B/n \\ L_{\rm tot} - \frac{\sigma_B\, A_{\rm tot}}{n}, & r > \sigma_B/n \end{cases} \quad \text{[S8a]}$$

$$B(t=\infty) = \begin{cases} \frac{L_{\rm tot}}{\sigma_B}, & r < \sigma_B/n \\ \frac{A_{\rm tot}}{n}, & r > \sigma_B/n \end{cases} \quad \text{[S8b]}$$

$$A_{\rm f}(t=\infty) = \begin{cases} A_{\rm tot} - \frac{n\, L_{\rm tot}}{\sigma_B}, & r < \sigma_B/n \\ 0, & r > \sigma_B/n\,. \end{cases} \quad \text{[S8c]}$$

Crucially, for $r < \sigma_B/n$, not all reactants are converted into products at the end of the reaction. Therefore, the $B(t = \infty)$ values increase proportionally with increasing $L_{\rm tot}$ values. These results are summarised in Supplementary Material Fig. S6. The value of the bifurcation point, $r^* = \sigma_B/n$, can be found by considering the steady states and conservation of binding sites, reactants and products in the system (where the transition in regimes occurs when $\theta_L = 0$ in the reactant limited regime) as follows:

$$\begin{aligned} \theta_L &= 1 - \theta_B \\ &= 1 - \frac{\sigma_B A_{\rm tot}}{n L_{\rm tot}}, \end{aligned} \quad \text{[S9]}$$

where we find:

$$r^* = \frac{\sigma_B}{n}\,. \quad \text{[S10]}$$

**S3.4. Early-time behavior.** Next, we examined the early-time dynamics of the system, as this allows the reaction term in Eq. (S7a) to be neglected as the rate of binding is significantly larger than the rate of conversion from $A$ to $B$. This means no conversion has taken place so $\theta_B = 0$ and Eq. (S7a) can be simplified as follows:

$$k_{\rm on}\left(A_{\rm tot} - \frac{L_{\rm tot}\theta_0}{\sigma_A}\right)(1-\theta_0) - k_{\rm off}\theta_0 = 0\,,$$

where $\theta_A(t \sim 0) = \theta_0$. This is rearranged to produce:

$$\frac{k_{\rm on}L_{\rm tot}}{\sigma_A}\theta_0^2 - \left(\frac{k_{\rm on}L_{\rm tot}}{\sigma_A} + k_{\rm on}A_{\rm tot} + k_{\rm off}\right)\theta_0 + k_{\rm on}A_{\rm tot} = 0\,,$$

which describes a quasi-instantaneous equilibrium formed by the binding and dissociation of A to the surface. The solution to this quadratic is:

$$\theta_0 = \frac{(\lambda + \rho\sigma_A + \sigma_A) - \sqrt{(\lambda + \rho\sigma_A + \sigma_A)^2 - 4\lambda\rho\sigma_A}}{2\lambda}\,, \quad \text{[S11]}$$

where:

$$\begin{aligned} \lambda &= \frac{L_{\rm tot}}{K_{\rm D}} \\ \rho &= \frac{A_{\rm tot}}{K_{\rm D}} \\ K_{\rm D} &= \frac{k_{\rm off}}{k_{\rm on}}\,. \end{aligned}$$

An equivalent form of Eq. (S11) is also used for convenience in later analysis:

$$\theta_0 = \frac{2\rho\sigma_A}{\lambda + \rho\sigma_A + \sigma_A + \sqrt{(\lambda + \rho\sigma_A + \sigma_A)^2 - 4\lambda\rho\sigma_A}} \,. \tag{S12}$$

Supplementary Material Fig. S7 compares the dynamics of $\theta_A$ and $\theta_B$, predicted by numerically evaluating the ODE system, with the early-time solution described in Eq. (S11). This demonstrates that the assumptions made in reaching Eq. (S11) are appropriate, as $\theta_A$ rapidly approaches $\theta_0$ before a relatively slow decline from this maximum as $A$ is converted to $B$ on the surface.

**S3.5. Half-time scaling.** To find analytical solutions, we use the early-time approximation where we set $\theta_A$ to be constant given by Eq. (S11). This allows Eq. (S7b) to be simplified to:

$$\frac{\mathrm{d}\theta_\mathrm{B}}{\mathrm{dt}} = nk_\mathrm{n}\theta_0^n \,, \tag{S13}$$

that has the solution (applying the initial condition: $\theta_B(t=0)=0$):

$$\theta_B(t) = nk_\mathrm{n}\theta_0^n t \,,$$

which can then be converted to describe the concentration of $B$:

$$B(t) = \frac{nk_\mathrm{n}L_\mathrm{tot}\theta_0^n t}{\sigma_B} \,. \tag{S14}$$

With an analytical solution describing $B(t)$, solutions for $t_{1/2}$ in the surface and reactant limited regimes, can be found by substituting $B(t=t_{1/2})$ in Eq. (S14).

- From the steady-state solutions, $B_{t_{1/2}}$ when $r < r^*$ (surface limited regime) is known to be $B_{t_{1/2}} = \frac{L_\mathrm{tot}}{2\sigma_B}$:

$$t_{1/2} = \frac{1}{2nk_n\theta_0^n} \,. \tag{S15}$$

- Again from the steady-state solutions, $B_{t_{1/2}}$ when $r > r^*$ (reactant limited regime) is known to be $B_{t_{1/2}} = \frac{A_\mathrm{tot}}{2n}$:

$$t_{1/2} = \frac{\sigma_B\rho}{2n^2k_n\lambda\theta_0^n} \,. \tag{S16}$$

To understand how Eq. (S11), Eq. (S15) and Eq. (S16) scale in the limit of $\rho$ or $\lambda$ approaching 0 or $\infty$, four cases were examined.

- **Case 1:** $\rho =$ constant & $\lambda \to 0$. In this limit the scaling of Eq. (S11) is examined first:

$$\lim_{\lambda\to 0}\theta_0 = \frac{\rho}{\rho+1} \,. \tag{S17}$$

  As $\lambda \to 0$, $r \to 0$, the system is in the surface limited regime and Eq. (S15) is used in finding the scaling of $t_{1/2}$ under these conditions, yielding:

$$t_{1/2} \propto \frac{1}{\theta_0^n} = \left(\frac{\rho+1}{\rho}\right)^n = \text{constant} \,, \tag{S18}$$

  since $\rho =$ constant.

- **Case 2:** $\rho =$ constant & $\lambda \to \infty$. From Eq. (S12):

$$\lim_{\lambda\to\infty}\theta_0 = \frac{\rho\sigma_A}{\lambda} \,. \tag{S19}$$

  Under these conditions, the system is in the reactant limited regime and Eq. (S16) scales as follows:

$$t_{1/2} \propto \frac{\rho}{\lambda\theta_0^n} = \sigma_A\left(\frac{\lambda}{\rho}\right)^{n-1} \,, \tag{S20}$$

  where $\rho$ and $\sigma_A$ are both constant so $t_{1/2}$ scales as:

$$t_{1/2} \propto \lambda^{n-1} \,. \tag{S21}$$

- **Case 3:** $\lambda = \text{constant}$ & $\rho \to 0$. From Eq. (S12):

$$\lim_{\rho \to 0} \theta_0 = \frac{\rho \sigma_A}{\lambda + \sigma_A}, \quad \text{[S22]}$$

and in the reactant limited regime, Eq. (S16) scales as:

$$t_{1/2} \propto \frac{\rho}{\lambda \theta_0^n} = \frac{\rho(\lambda + \sigma_A)^n}{\lambda(\rho\sigma_A)^n}, \quad \text{[S23]}$$

and therefore

$$t_{1/2} \propto \rho^{-(n-1)}. \quad \text{[S24]}$$

- **Case 4:** $\lambda = \text{constant}$ & $\rho \to \infty$. From Eq. (S12):

$$\lim_{\rho \to \infty} \theta_0 = 1, \quad \text{[S25]}$$

and in the surface limited regime, Eq. (S15) scales as:

$$t_{1/2} \propto \frac{1}{\theta_0^n} = \text{constant}. \quad \text{[S26]}$$

These scaling results can be summarized by considering $\lambda \to \infty$ and $\rho \to 0$ as equivalent to describing $r \to \infty$. Similarly, $\lambda \to 0$ and $\rho \to \infty$ describe $r \to 0$. The results from the four different scaling conditions can be converted to be in terms of $r$:

$$t_{1/2} \sim r^{\gamma},$$

where the scaling exponent, $\gamma$, displays biphasic behavior depending on $r$. The analytical expression describing $t_{1/2}$ for the reactant limited regime, Eq. (S16), of the system's behavior has a stationary point, $r_s$, that is found by taking the derivative with respect to $L_{\text{tot}}$, producing:

$$r_s = \frac{K_{\text{D}}}{A_{\text{tot}}} + 1,$$

where we assumed $n = 2$ for this specific result. The scaling analysis is summarized in Supplementary Material Fig. S8 where the intermediate regime between the surface and reactant limited regimes is defined as $r^* < r < r_s$ and the $t_{1/2}$ behavior does not obey the scaling predictions.

**S3.6. Self-consistent analytical solution.** The first step in deriving the analytical solution is using an asymptotic approach assuming that we can separate the binding-dissociation kinetics from the much slower chemical reaction. Under these conditions, the binding pre-equilibrium amounts to setting the binding term in the kinetic equations Eq. (S7) to zero:

$$0 = k_{\text{on}} \left( m_{\text{tot}} - L_{\text{tot}} \left[ \frac{\theta_A}{\sigma_A} + \frac{\theta_B}{\sigma_B} \right] \right) (1 - \theta_A - \theta_B) - k_{\text{off}}\, \theta_A. \quad \text{[S27]}$$

This equation can be solved for $\theta_A$ as a function of $\theta_B$, which can be approximated as a linear function using the values of $\theta_A(\theta_B) = 0$ and $\theta_A(\theta_B = 0)$ as reference points. Setting $\theta_B = 0$ in the pre-equilibrium Eq. (S27) gives the same result as the early-time solution Eq. (S11) $\theta_A(t = 0) = \theta_0$, while setting $\theta_A = 0$ amounts to computing the steady-state solution of $\theta_B$ and has two solutions $\theta_B = 1$ and $\theta_B = \frac{\sigma_B}{r}$ with $r = \frac{L_{\text{tot}}}{A_{\text{tot}}}$. As $\theta_B$ cannot exceed 1, the second solution is only physically valid for $r > \sigma_B/n$. This allows us to derive the linearized dependence of $\theta_A$ on $\theta_B$ to be:

$$\theta_A(\theta_B) = \theta_0 \left(1 - \frac{\theta_B}{A}\right)$$
$$A = \begin{cases} 1 & , r < \sigma_B \\ \frac{\sigma_B}{r} & , r \geq \sigma_B, \end{cases} \quad \text{[S28]}$$

which can be used to reduce the number of kinetic equations by one and hence facilitates solving the equations. Plugging the above expression into the kinetic equation for $\theta_B(t)$ yields:

$$\frac{d\theta_B}{dt} = k_n\,\theta_0^n \left(1 - \frac{\theta_B}{A}\right)^n \tag{S29}$$

which can be solved by integration resulting in:

$$\theta_B(t) = \begin{cases} A\left(1 - \left(1 - \frac{\theta_B(0)}{A}\right) e^{-\frac{k_n \theta_0}{A} t}\right), & \text{for } n = 1 \\ A\left(1 - \left(\frac{(n-1)\,k_n\,\theta_0^n}{A}\,t + \left(1 - \frac{\theta_B(0)}{A}\right)^{1-n}\right)^{\frac{1}{1-n}}\right), & \text{for } n > 1. \end{cases} \tag{S30}$$

The plateau value of $\theta_B(t)$ can readily computed by taking the limit $t \to \infty$ yielding:

$$\theta_B(t \to \infty) = A\,, \tag{S31}$$

for arbitrary $n$ and any initial value $\theta_B(0)$. The solution for $\theta_A(t)$ is then given by plugging the solution for $\theta_B(t)$ back into Eq. (S28). The half-time scaling can then be computed by solving $\theta_B(t_{1/2}) = \frac{1}{2}$ yielding:

$$t_{1/2} = \begin{cases} \frac{A}{k_n\,\theta_0} \log\left(\frac{2(A-\theta_B(0))}{A}\right), & \text{for } n = 1 \\ \frac{A}{(n-1)\,k_n\,\theta_0^n}\left(\left(\frac{1}{2}\right)^{1-n} - \left(1 - \frac{\theta_B(0)}{A}\right)^{1-n}\right), & \text{for } n > 1. \end{cases} \tag{S32}$$

### S4. Low lipid-to-protein ratios

At low lipid-to-protein ratios, $r < 2$, the experimental kinetic traces exhibit the key qualitative feature of the lipid-limited ($2 < r < \alpha$) regime, a lipid-dependent yield, but also have a lipid-dependent kinetic profile. This lipid-dependent kinetic profile is displayed in Supplementary Material Fig. S9, where the kinetic traces slow down for decreasing $r$ values analogous to the behavior in the protein-limited regime ($r > \alpha$) driven by the dilution effect on the surface coverage of protein monomers. Previous investigations into what drives this change in behavior at low $r$ values concluded that lipid-induced elongation becomes a two-step process due to the low abundance of lipids (2). As our model implicitly assumes the lipid-dependent elongation process is saturated with respect to lipids for the experimental conditions considered in this study, our theoretical model used to analyze the data presented in Fig. 3 cannot accurately fit the experimental data at low $r$ values (Supplementary Material Fig. S9a).

To address this limitation, we can modify the reaction flux of nucleation of fibrils in the one-step primary nucleation model as follows:

$$J_{\text{nucleation}} = \frac{k_{\text{n}}}{L_{\text{tot}}} \left( \frac{\theta_m}{1 + \frac{r^*}{r}} \right)^{n_1} \left( m_{\text{tot}} - L_{\text{tot}} \left[ \frac{\theta_m}{\beta} + \frac{\theta_M}{\alpha} \right] \right)^{n_2} , \qquad \text{[S33]}$$

where $r^*$ is the critical lipid-to-protein ratio below which we observe lipid-dependent kinetic profiles. Previous work indicates $r^* = 2$ and this value is used to fit to the experimental data in Supplementary Material Fig. S9 (2). The reaction flux of nucleation of oligomers in the two-step primary nucleation model is similarly modified to:

$$J_{\text{nucleation}} = k_{\text{o}} \left( \frac{\theta_m}{1 + \frac{r^*}{r}} \right)^{n_1} \left( m_{\text{tot}} - L_{\text{tot}} \left[ \frac{\theta_m}{\beta} + \frac{\theta_M}{\alpha} \right] \right)^{n_2} , \qquad \text{[S34]}$$

and finally, the reaction flux of elongation for one-step and two-step models is modified to:

$$J_{\text{elongation}} = 2k_+ \left( \frac{\theta_m}{1 + \frac{r^*}{r}} \right) \theta_P \, . \qquad \text{[S35]}$$

The addition of the $1/(1+r^*/r)$ term can be intuitively interpreted as follows. When $r < r^*$, the term $r^\star/r$ dominates, and thus:

$$\frac{\theta_m}{1 + \frac{r^*}{r}} \sim \frac{\theta_m \, r}{r^\star} \, .$$

Since $\theta_m = \beta \, m_{\text{b}}/L_{\text{tot}}$, this reveals that the lipid-dependence for the relevant aggregation processes at $r < r^\star$ is determined by:

$$\theta_m \, r = \frac{\beta \, m_{\text{b}}}{m_{\text{tot}}} \, ,$$

where $m_{\text{b}}$ is limited physically by the availability of free lipid binding sites. Alternatively, if $r > r^\star$

$$\frac{\theta_m}{1 + \frac{r^*}{r}} \sim \theta_m \, ,$$

where the extension recovers the original form of the model. The ability to quantitatively describe the experimental data at low lipid-to-protein ratios is demonstrated through the fits summarized in Supplementary Material Fig. S9b. Details of the fitting result can be found in Supplementary Material Table S10. This addition to the model changes the analytical solution simply as follows:

$$F^*(t) = F(t) \left( \frac{1}{1 + \frac{r^*}{r}} \right)^{n_1 + 1} , \qquad \text{[S36]}$$

where $F^*(t)$ is the adjusted $F(t)$ function, with various definitions for the models shown in the Methods section.

### S5. Cooperative binding of protein monomer to lipid surface

In our model of lipid-induced $\alpha$-synuclein aggregation, we have assumed non-cooperative binding of the $\alpha$-synuclein monomers to the lipid surface. Cooperative binding is where the probability of a protein monomer binding to a given surface site depends on the binding state of neighboring sites. Strong cooperativity, with binding occurring favorably at sites where neighboring sites are also bound by other protein monomers, results in individual vesicles being covered completely by bound protein before another vesicle begins to be occupied.

To account for a cooperative binding process, the description of the reaction flux due to protein-lipid binding can be extended to:

$$J_{\text{binding}} = k_{\text{on}} \left( m_{\text{tot}} - L_{\text{tot}} \left[ \frac{\theta_m}{\beta} + \frac{\theta_M}{\alpha} \right] \right)^{n_{\text{b}}} (1 - \theta_m - \theta_M) - k_{\text{off}}\, \theta_m \,, \qquad \text{[S37]}$$

where $n_{\text{b}}$ is analogous to the Hill coefficient in the Hill Equation for describing cooperative binding. The key behaviors of this model are summarized in Supplementary Material Fig. S10. There is no effect on the system's steady state behavior. The predicted kinetics are quantitatively affected, which can be compensated by adjusting the reaction rates, but the critical qualitative features of the system's kinetic behavior are unchanged.

There is strong evidence that $\alpha$-synuclein binds cooperatively to some lipid surfaces (4). However, there is no substantive evidence on whether the binding is cooperative for the lipid-induced aggregation system that is the focus of this work. For this reason, along with the qualitative similarities demonstrated in Supplementary Material Fig. S10, cooperative binding was neglected from the model and subsequent analysis of experimental data for the lipid-induced aggregation kinetics.
